# Structure of trimeric pre-fusion rabies virus glycoprotein in complex with two protective antibodies

**DOI:** 10.1101/2022.02.28.482293

**Authors:** Weng M Ng, Sofiya Fedosyuk, Solomon English, Gilles Augusto, Adam Berg, Luke Thorley, Rameswara R Segireddy, Thomas A Bowden, Alexander D Douglas

## Abstract

Rabies virus (RABV) causes lethal encephalitis and is responsible for approximately 60,000 deaths per year. As the sole virion-surface protein, the rabies virus glycoprotein (RABV-G) mediates host-cell entry. RABV-G’s pre-fusion trimeric conformation displays epitopes bound by protective neutralizing antibodies which can be induced by vaccination or passively administered for post-exposure prophylaxis. We report a 2.8-Å structure of a RABV-G trimer in the pre-fusion conformation, in complex with two neutralizing and protective monoclonal antibodies, 17C7 and 1112-1. One of these antibodies is a licensed prophylactic (17C7, Rabishield), which we show locks the protein in pre-fusion conformation. We demonstrate that targeted mutations can stabilize RABV-G in the pre-fusion conformation, a key step towards structure-guided vaccine design. These data reveal the higher-order architecture of a key therapeutic target and the structural basis of neutralization by antibodies binding two key antigenic sites, and will facilitate the development of improved vaccines and prophylactic antibodies.

## Introduction

An estimated 3 billion people live at risk of rabies virus (RABV; genus *Lyssavirus* and family *Rhabdoviridae*) infection, which causes fatal encephalitis (Fooks et al., 2017; Hampson et al., 2015). Although effective pre- and post-exposure vaccines and passive immunization treatments are available, their high cost and need for multiple doses to achieve protection result in inadequate coverage of at-risk populations (2018).

The RABV envelope surface displays the virion glycoprotein RABV-G, a trimeric class III viral fusion protein, which mediates receptor binding and membrane fusion during host-cell entry. Multiple host proteins have been implicated in RABV cell entry in different contexts, but no single RABV-G : receptor interaction has been shown to be indispensable across contexts (Lentz et al., 1982; Thoulouze et al., 1998; Tuffereau et al., 2007; Wang et al., 2018). In contrast to class I viral fusion proteins, which are also trimeric and are better-studied, RABV-G can transition reversibly between a pre-fusion form (predominant at neutral pH) and post-fusion form (predominant at acidic pH) (Gaudin et al., 1992; Gaudin et al., 1991).

As the sole virion surface protein, RABV-G is the primary target of protective antibodies (Gaudin et al., 1999; Wiktor et al., 1973). Historically, polyclonal rabies immune globulin (RIG) has been used for post-exposure passive immunization. The use of anti-RABV-G neutralizing monoclonal antibodies (mAbs) is now attracting increasing attention as an alternative to expensive and often human-donor-derived rabies immune globulin (Sparrow et al., 2019). Two anti-rabies mAb products have been licensed for clinical use in India, which experiences more rabies cases than any other single country: one is a single antibody (Rabishield™, 17C7 or RAB1 (Gogtay et al., 2018; Sloan et al., 2007)); the other is a combination of two mAbs (Twinrab™, docaravimab/miromavimab, also known as 62-71-3 and M777 (Muller et al., 2009)). Draft guidance to industry regarding the path to potential approval of such mAbs for the US market has recently been published by the US FDA (FDA, 2021). A key consideration in the development of such therapies is the breadth of coverage against circulating rabies isolates, and ideally also against related bat lyssaviruses, which also cause human disease. In this regard, there is some concern about the vulnerability of 17C7, administered as a single mAb, to known antigenic polymorphisms (De Benedictis et al., 2016; Sloan et al., 2007).

Development of subunit protein and mRNA-based rabies vaccines has, so far, proven challenging (Aldrich et al., 2021; Ertl, 2019). It is believed that the trimeric pre-fusion form is the ideal immunogen, as its surface displays the major known neutralizing antibody (nAb) epitopes and monomeric proteins have performed poorly as immunogens (De Benedictis et al., 2016; Koraka et al., 2014). To our knowledge, however, production of soluble and stable pre-fusion trimeric recombinant RABV-G has not been reported. Previous efforts to structurally characterize RABV-G produced crystal structures of a single domain and monomeric ectodomains of RABV-G in complex with nAbs, verifying that RABV-G is a class III fusion glycoprotein (Hellert et al., 2020; Yang et al., 2020). Extensive efforts over decades have, however, been unsuccessful in obtaining high resolution insight into either the protein’s higher-order architecture or complexes of nAbs with the biologically and antigenically critical pre-fusion conformation. This information would empower the rational design of improved RABV vaccine immunogens and antibody-based therapeutic cocktails.

## Results

### Structure of trimeric pre-fusion RABV-G

To characterize trimeric RABV-G, we recombinantly expressed a construct of the full-length G trimer (Lys1–Leu505) encoding a site-directed mutation (H270P), which we designed with the intention of stabilizing the pre-fusion conformation. His270 lies within an elongated alpha-helix in a previous crystal structure of monomeric RABV-G ectodomain (RABV-G^ecto^) at low pH (likely post-fusion conformation), and a pre-fusion stabilizing effect of the analogous L271P mutation upon the glycoprotein of another rhabdovirus (vesicular stomatitis virus, VSV) has previously been reported (Ferlin et al., 2014; Yang et al., 2020). Detergent-extracted RABV-G was complexed with the antigen-binding fragments (Fab) of mAb 17C7 and mAb 1112-1 and subjected to single-particle cryo-electron microscopy (cryo-EM) analysis to generate a 2.8-Å resolution structure of a RABV-G trimer – Fab 17C7 – Fab 1112-1 complex (Fig. 1, Supplementary Fig. 1–2, and Supplementary Table 1). As observed for other class III fusogens (Roche et al., 2006), each protomer of the trimer is composed of three domains: a membrane-distal pleckstrin homology domain (PHD), a membrane-proximal fusion domain (FD), and a laterally positioned central domain (CD) (Fig. 1D–E). Topologically, PHD sits atop the FD, and the CD is located adjacent to the PHD/FD junction. The PHD is connected to the CD via two linkers Leu22–Asn37 (termed L1) and Gly255–Leu271 (L4), respectively, and to the FD via two separate linkers consisting of residues Lys47–Ser52 (L2) and Glu181–Ile191 (L3). The fusion loops, transmembrane, and intraviral regions were not resolved in the reconstruction, indicative that these regions of the molecule exhibit elevated levels of flexibility in the purified protein.

**Figure 1:**
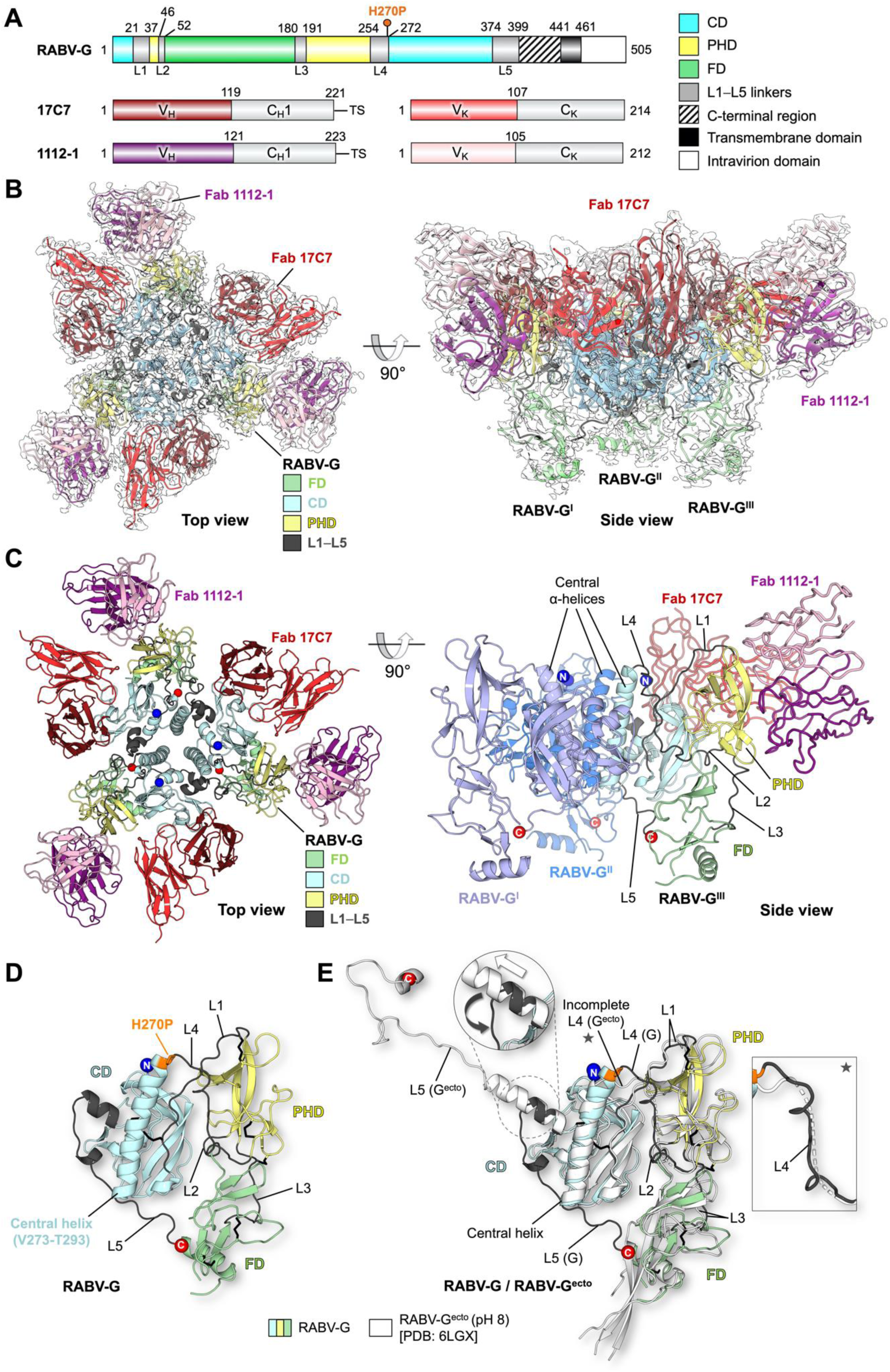
Structure of pre-fusion RABV-G trimer in complex with Fabs 17C7 and 1112-1. **(A) Schematic of RABV-G domain boundaries** Linear map of RABV-G protein sequence is drawn to scale using DOG software (Ren et al., 2009), with domains colored as indicated in the legend, showing ‘palindromic’ architecture typical of class III fusion proteins (CD: central domain; PHD: pleckstrin homology domain; FD: fusion domain). The H270P point mutation in our protein construct is indicated with a pin above the map. C-terminal region (shaded) is unresolved in our structure. (*Bottom*) Linear maps of Fabs 17C7 and 1112-1 heavy and kappa light chains, colored and labeled accordingly. V_H_, V_K_, C_H1_, and C_K_ denote the antibody variable heavy, variable kappa light, constant 1 heavy, and constant kappa light-chain domains, respectively. TS, TwinStrep tag. **(B) 2.8-Å cryo-EM map with the resulting structure of trimeric RABV-G shown in top and side view orientations**. Single copies of 17C7 and 1112-1 were observed to bind to each protomer of RABV-G. The atomic model is fitted into the corresponding cryo-EM map (white) and colored according to domain with the variable regions of Fab 17C7 and Fab 1112-1 colored red and purple (darker shade for heavy chain, lighter shade for light chain), respectively. The constant regions of the Fabs were disordered in the reconstruction and therefore were not built. **(C) Atomic model of the RABV-G – Fab 17C7 – Fab 1112-1 complex**. (*Top view*) The protein molecules are displayed in cartoon representation and colored accordingly as labeled. N- and C-termini are shown as blue and red spheres, respectively. (*Side view*) Only one copy of each Fab is shown in ribbon representation. Two RABV-G protomers are colored blue and light purple for visual clarity. The remaining copy is colored as shown in top view. **(D) Conformational features revealed by atomic model of RABV-G**. A single protomer of RABV-G is shown in cartoon representation. PHD is colored yellow, CD in blue, and FD in green, while the inter-domain linkers (L1–L5) are colored in dark gray. N- and C-termini are shown as blue and red spheres, respectively. The point mutation H270P is colored orange and shown in stick representation. **(E) Structure superimposition of our trimeric RABV-G with a previously reported monomeric RABV-G ectodomain structure obtained at pH 8.0** (RABV-G^ecto^, white cartoon, PDB ID: 6LGX) (Yang et al., 2020). When domains were aligned separately, the PHD and CD aligned more closely than the FD (calculated root-mean-square deviations 1.0-Å over 72 equivalent Cα atoms for PHD, and 0.6-Å/128 Cα for CD, and 2.4-Å/88 Cα for FD). Differences in the L4 and L5 linkers between our trimeric RABV-G structure and the previous RABV-G^ecto^ structure are highlighted in the inset: additional details regarding these linkers are provided in Supplementary Fig. 3–4.

Structural overlay of our trimeric RABV-G with the previously reported monomeric RABV-G^ecto^ determined at pH 8.0 reveals good structural agreement between individual domains, but significant dissimilarity in inter-domain linkers, rearrangement of which is believed to be critical for the pre-to post-fusion conformational transition (Fig. 1E) (Roche et al., 2007). The region from Pro374 to the C-terminus in the high-pH crystal structure of G^ecto^ forms a helical structure distal from the FD. We term the equivalent region of our trimeric RABV-G ‘L5’, extending from Pro374 to the C-terminal limit of our model at Leu399. L5 forms a loop which interacts with the adjacent protomer’s CD and then runs back across the base of the CD to reach the FD (Supplementary Fig. 3). The interaction interface between the L5 loop and the FD contains several hydrophobic interactions, including contributions by a cluster of histidines (His86, His173 and His397; Supplementary Fig. 3B). As has been suggested for a similar histidine cluster in VSV-G (Roche et al., 2007), protonation of these residues at low pH may act as a ‘pH sensor’, disrupting the L5–FD interface and initiating the movement of FD towards the target membrane. The different orientations of the L5 region in our RABV-G and RABV-G^ecto^ are similar to the pre-fusion and intermediate conformations of VSV-G and Chandipura virus G (CHAV-G) (Baquero et al., 2012; Roche et al., 2007), respectively. It seems possible that the RABV-G^ecto^ structure, which lacks membrane interactions and is potentially influenced by the packing environment of the protein crystal, may represent such an intermediate.

A second notable difference between trimeric RABV-G and the RABV-G^ecto^ pH 8.0 crystal structure lies in linker L4, which connects the central helix of the CD with the PHD. While the linker was not fully observed in the G^ecto^ structure, it is resolved in our structure due to the stabilizing contacts mediated by the trimeric organization of the molecule (Fig. 1E and Supplementary Fig. 4). This region forms an α-helix in the previously reported low-pH RABV-G^ecto^ structure and contains both the helix-breaking H270P substitution introduced here and a series of further amino acids, which have previously been implicated in the fusion competence of the protein (His261, Asp266 and Glu269) (Yang et al., 2020). Conservation of the local architecture in the region of Pro270 in our structure, as compared to the pre-fusion VSV-G structure and high pH RABV-G^ecto^ crystal structure (Fig. 1E and Supplementary Fig. 5) (Roche et al., 2007; Yang et al., 2020), suggests that this region of our protein retains an authentic conformation. We observe that L4 mediates a series of hydrophobic contacts with the PHD and FD, and hydrogen bonds with L1 and the central helix (Supplementary Fig. 4). Separation of these contacts would be necessary to allow hinging movement of the PHD relative to the CD, which is thought to be required during rearrangement of class III proteins into post-fusion conformation (Roche et al., 2007). This may be facilitated by protonation of His261, disrupting the pre-fusion hydrogen bond with Asn26 in L1, and allowing formation of the interaction with Asp211 as previously visualized in RABV-G^ecto^ low pH structure (Yang et al., 2020).

The conformations of L4 and L5 revealed by our structure thus give clues about the roles these regions play during the fusogenic conformational change of RABV-G, where dislocation of the L5 loop from the FD and separation of the L4 from the PHD are required to create an extended intermediate. This is consistent with a model based upon low-pH structures of VSV-G, CHAV-G and RABV-G, whereby L4 residues from 266 onwards then extend the central helix, with an adjacent antiparallel helix formed by re-folding of L5 (Albertini et al., 2012; Baquero et al., 2015; Yang et al., 2020).

To further explore the role of L4 and L5 in pH-mediated conformational transition, we used a flow cytometry-based assay to assess the effect of targeted mutations on conformational stability of RABV-G (Fig. 2). Of 12 targeted histidine loci across the protein, H261A/L mutations (in L4) had the greatest stabilising effect (a 3-fold increase in retention of the pre-fusion conformation at pH 5.8). These mutations would prevent the electrostatic interaction observed between His261 and Asp211 in the low pH G^ecto^ structure, and their effect suggests that the His261-Asn26 hydrogen bond is dispensable for the pre-fusion structure. Attempted mutations at the L5–FD interface proved incompatible with protein expression (Supplementary Table 2). We also assessed the effect of substitution with proline of each residue from Iso268 to Val272 (in L4), and of His384 (in L5). On the basis of comparison to previous low pH RABV-G^ecto^, CHAV-G and VSV-G structures (Baquero et al., 2015; Roche et al., 2006; Yang et al., 2020), these residues are likely to lie in extended helices in the post-fusion conformation, formation of which may be disrupted by proline. H270P substitution had a marked stabilising effect (3 to 4-fold increase in retention of pre-fusion conformation at pH 5.8) (Fig. 2B). H270P and H261A/L preserved approximately wild-type expression levels and antigenicity (Fig. 2C and Supplementary Table 2). These represent important steps towards structure-guided design of stable RABV-G subunit vaccines, and the information we provide regarding L4 and L5 structure should facilitate design of improved future constructs.

**Figure 2:**
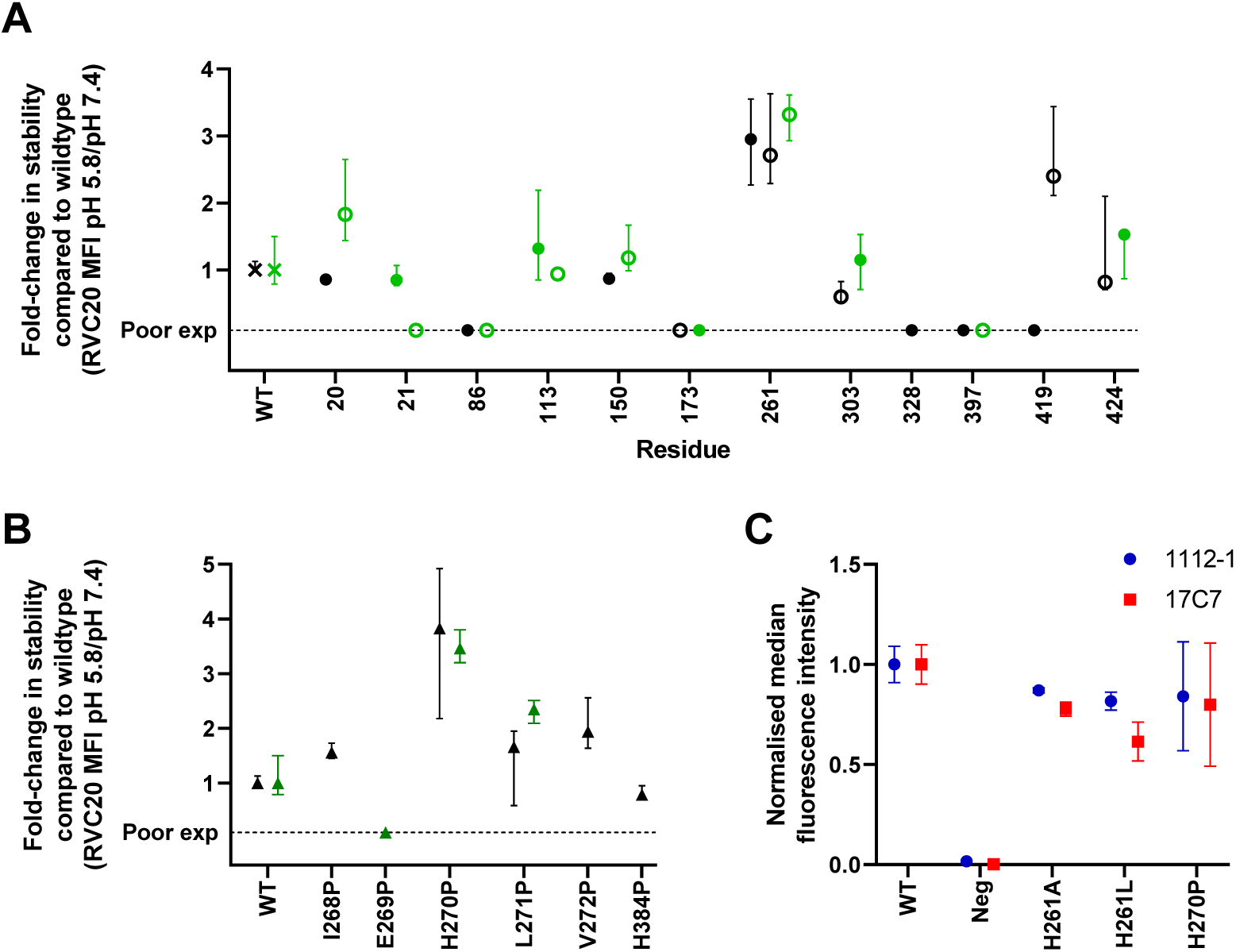
Targeted mutations at two sites in L4 stabilize pre-fusion RABV-G. Wildtype (WT) and mutant RABV-G constructs were expressed on transiently transfected Expi293 cells, and reactivity with site I (RVC20), site II (1112-1) and site III (17C7) IgG was assessed by flow cytometry. Expression levels of all constructs are shown in Supplementary Table 2. **(A) Effect of histidine substitutions**. All histidine residues in the RABV-G ectodomain were mutated to alanine and to leucine, with exceptions detailed in methods. Pre-fusion protein stability was measured by calculating median fluorescence intensity (MFI) of the pre-fusion-specific mAb RVC20 (De Benedictis et al., 2016; Hellert et al., 2020) after binding at pH 5.8 as a proportion of that after binding to the same construct at pH 7.4: this proportion was 0.11 for untagged WT RABV-G, and 0.12 for WT RABV-G expressed as a fusion protein. Results are expressed as fold-change in this proportion as compared to WT protein. Filled and open symbols denote introduction of alanine and leucine respectively. Black and green symbols denote untagged constructs and those expressed as GFP fusion proteins respectively. Points represent median and error bars represent range of four technical replicates across two experiments (a transfection with each of two independent DNA preparations on each of two days). ‘Poor exp’ denotes constructs which expressed at <33% of the level of WT RABV-G, as assessed by RVC20 binding at pH 7.4 **(B) Effect of potentially helix-breaking substitutions with proline**. Residues in L4 / L5 regions expected to form helices in post-fusion protein were substituted with proline. Colours, replication (points and error bars) and the definition of poor expression are as for panel (A). **(C) H261A/L and H270P retain site II and site III antigenicity, as evidenced by 1112-1 and 17C7 binding**. Untagged constructs were used. MFI is expressed as a proportion of that observed with WT RABV-G with each antibody. ‘Neg’ denotes MFI on cells transfected with an irrelevant antigen. The replication strategy and meaning of points and error bars were as for panel (A).

### Trimerization of RABV-G

In our structure, RABV-G monomers assemble as a compact tripod with the CDs located adjacent to the trimerization axis (Fig. 3). Each CD presents a near-vertically oriented α-helix (Val273–Thr293) that packs and forms the trimerization core. Similar to VSV-G, the extent of the inter-protomeric interface (∼1,600 Å^2^) is approximately 1/7 of that in a pre-fusion herpesvirus class III fusion protein, human cytomegalovirus gB (∼11,200 Å^2^).

**Figure 3:**
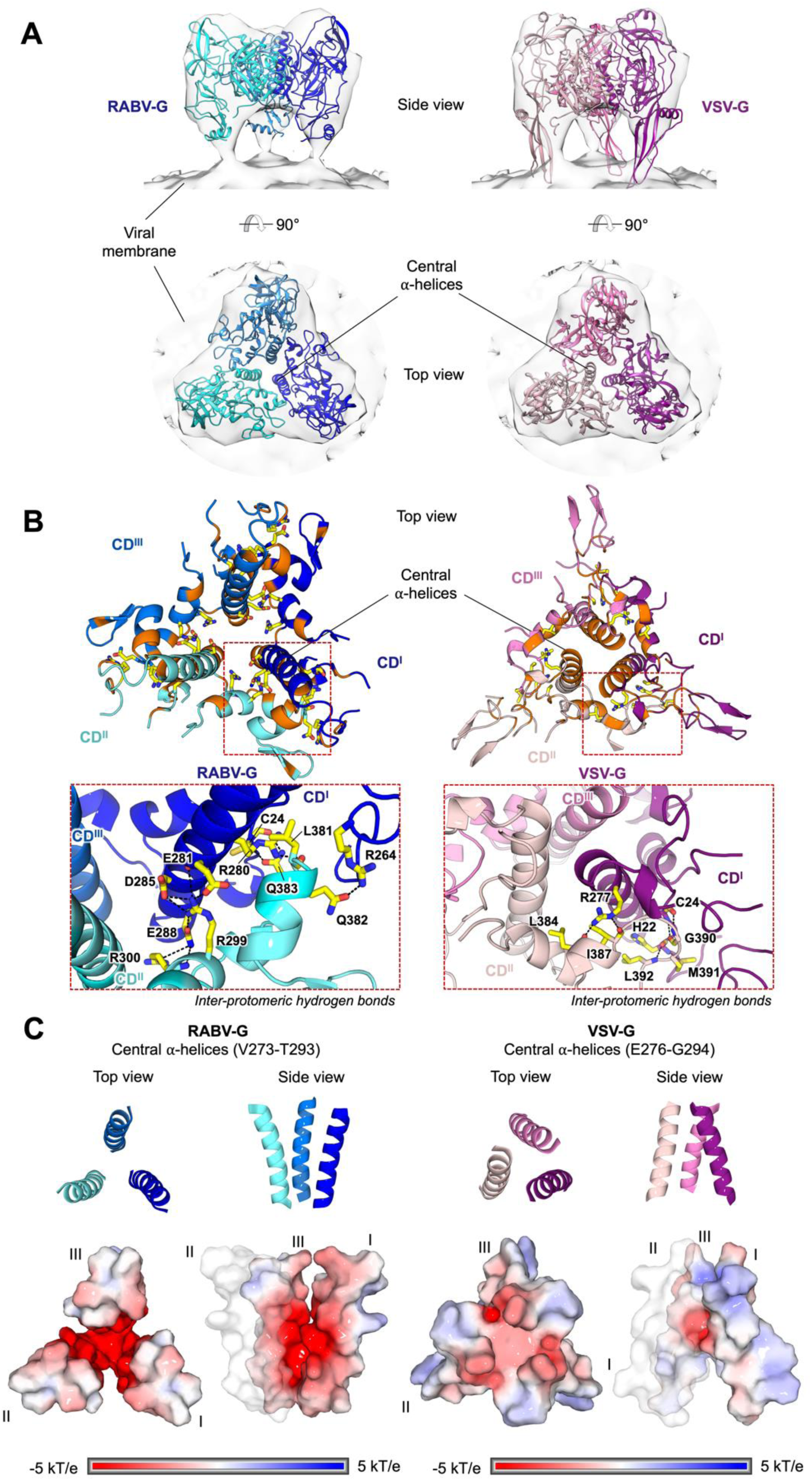
Structural comparison of pre-fusion RABV-G and VSV-G reveals contrasting modes of inter-protomeric interactions at the trimerization core. **(A) Fitting of pre-fusion RABV-G (blue) and VSV-G (pink; PDB ID: 5I2S) into a subtomographic average map of pre-fusion VSV-G (semi-transparent gray; EMD-9331**) (Si et al., 2018). RABV-G and VSV-G structures are shown in cartoon representation and colored in different shades of blue and pink, respectively. Maps are shown as transparent surfaces. **(B) Zoom-in views of the G trimerization core involved in inter-protomeric interactions**. Residues involved in hydrogen bonding (black dashed lines) are shown as sticks, with the carbon, nitrogen, oxygen, and sulfur constituents colored yellow, blue, red, and dark yellow, respectively. Residues involved in non-polar interactions are colored orange. The zoom-in panels of the CD α-helices shows the inter-protomeric hydrogen bonds. This analysis demonstrates that the trimerization core of RABV-G is largely mediated by polar interactions between charged residues, including a network of hydrogen bonds formed by negatively-charged Glu281, Asp285, Glu288 on one protomer and positively-charged Arg299 and Arg300 on the adjacent protomer. Hydrophobic interactions are formed at the periphery and bottom of the central α-helices. In contrast, VSV-G displays the reverse pattern of inter-protomeric interactions, where the core is largely maintained by hydrophobic interactions with hydrogen bonds formed at the periphery. **(C) Electrostatic potential of the central α-helices**. The central α-helices of RABV-G (left) and VSV-G (right) are shown as cartoons (top) and surfaces (bottom). The surfaces are colored according to the electrostatic potential in the range of ± 5 kT/e, as calculated by Adaptive Poisson-Boltzmann Solver (APBS). Both RABV-G and VSV-G display negatively-charged trimerization cores. This characteristic is especially prominent in RABV-G, where the carboxyl groups of Glu274, Glu275, Glu281, Glu282, Asp285, and Glu288 side chains line the central α-helices.

Other features of the RABV-G trimerization interface, however, contrast with the pre-fusion VSV-G structure. Notably, the central helices in our RABV-G structure form an inverted cone with the apex proximal to the membrane, while in VSV-G the apex of the analogous cone is distal to the membrane (Fig. 3C). This reflects differing relative angulation of the entire protomers between the two structures (i.e. the orientation of the RABV-G central helix relative to the remainder of the RABV-G protomer in our structure is similar to that seen in VSV-G) (Fig. 3A and Supplementary Fig. 5). In addition to the contributions of the L4 and L5 loops to intra-protomeric interactions (Supplementary Fig. 3–4), as described above, the inter-protomeric interactions include hydrogen bonds formed by Leu381, Gln382, and Gln383 (within L5) with Arg280 (within central helix), Arg264 (within L4), and Cys24 (within L1), respectively (Fig. 3B). Transient dissociation of the RABV-G trimer is believed to be required to permit rearrangement to the post-fusion conformation (Albertini et al., 2012): the fact that the inner faces of RABV-G’s central helices are lined by acidic residues and hence markedly electronegative may facilitate this (Fig. 3C). The information we provide regarding the RABV-G trimer interface, which has not previously been characterized, should facilitate design of constructs to enhance the stability of the trimer.

### Structural basis of RABV-G binding by protective antibodies

Previous antibody epitope mapping studies have revealed three major antigenic sites on RABV-G, designated I–III, and two less-commonly recognized sites, IV and ‘a’ (Benmansour et al., 1991; Kuzmina et al., 2013) (Supplementary Fig. 6). Here, we reveal the molecular specificity underlying antibody-mediated targeting by the Fabs of two mAbs, 1112-1 and 17C7, which are both protective against RABV challenge in animal models and bind to sites II and III respectively (Dietzschold et al., 1992; Sloan et al., 2007).

In our structure, Fabs 17C7 and 1112-1 bind to each protomer of the RABV-G trimer (Fig. 4). Fab 17C7 binds CD with an interaction interface nearly parallel to the membrane, whilst Fab 1112-1 engages the PHD with an interface nearly perpendicular to the membrane. This alternating longitudinal and latitudinal binding to adjacent epitopes brings the two Fabs in close proximity, such that contacts are observed between the 17C7 light chain and the 1112-1 heavy and light chains. No ordered density corresponding to glycans was seen in our map (Supplementary Fig. 7), but the 1112-1 epitope is adjacent to the potential glycosylation sites Asn37 (believed to have low glycan occupancy (Yamada et al., 2013)) and Asn247.

**Fig. 4.**
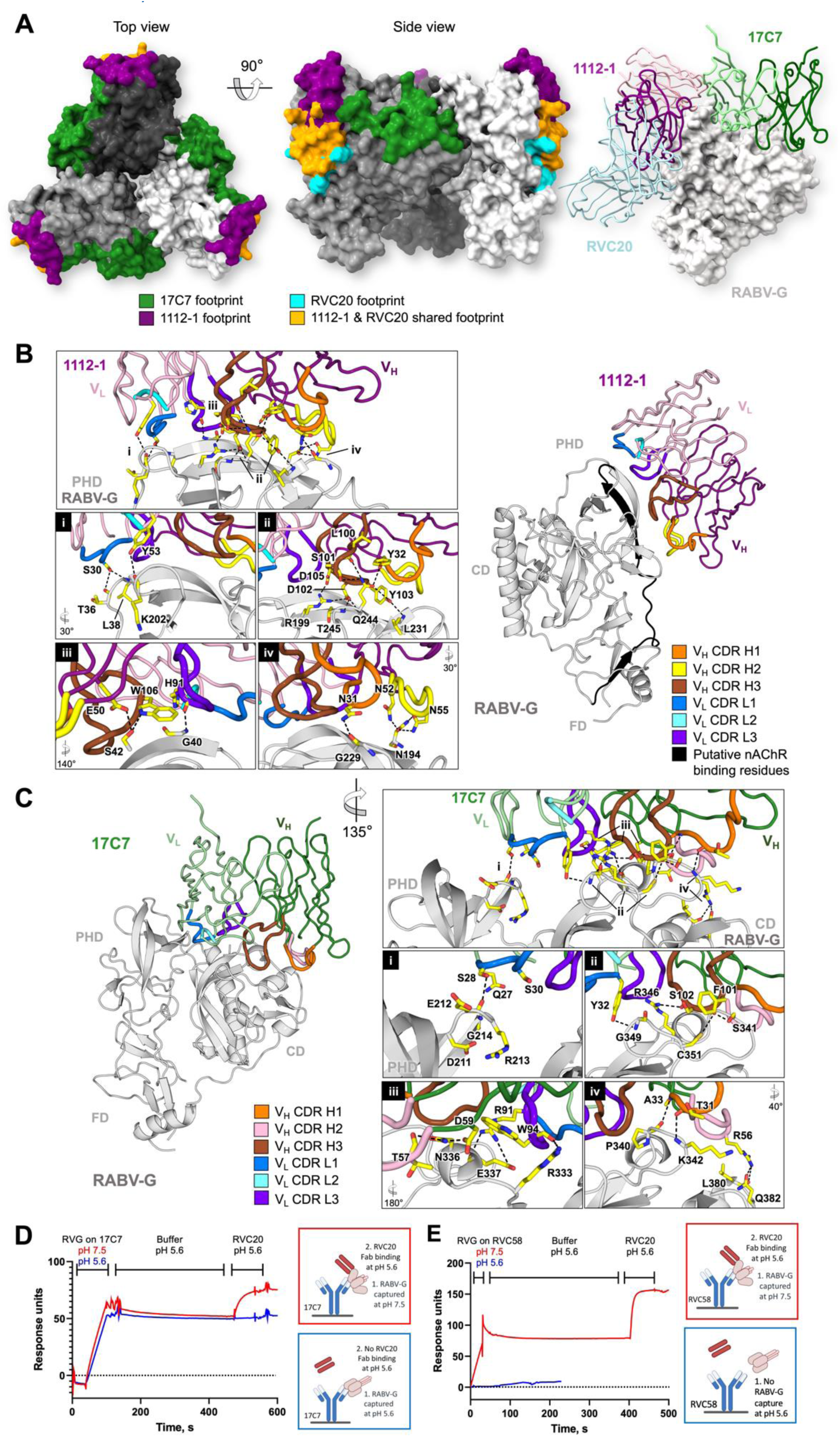
Structural basis for antibody-mediated RABV neutralization: multiple sites of vulnerability on the membrane-distal crown of the RABV-G trimer. **(A) Visualization of structurally characterized anti-RABV antibody epitopes**. (*Left*) Footprints exhibited by RVC20 (site I-targeting antibody, PDB ID: 6TOU; cyan), 1112-1 (site II; purple), and 17C7 (site III; green) plotted on trimeric RABV-G (white, gray, and dark gray surface). MAbs 1112-1 and RVC20 are shown to target distinct yet overlapping epitopes (orange surface). For clarity, the variable regions of the Fab fragments of mAbs RVC20, 1112-1 and 17C7 bound to a single RABV-G protomer are shown as ribbons (*right*). **MAbs (B) 1112-1 and (C) 17C7 target the PHD and CD domains of RABV-G, respectively**. Detailed interactions between complementarity-determining regions (CDRs) of each antibody with RABV-G (gray cartoon) are highlighted in the boxed panels. Residues involved in the antibody–antigen interactions are shown as yellow sticks, with the CDR loops colored as indicated. In our structure, one molecule of each Fab binds to each protomer of the RABV-G trimer. **(B)** The 1112-1 epitope encompasses three β-strands and five loops on the PHD, which include residues 175-203 (highlighted black) that have been previously implicated in nicotinic acetylcholine receptor (nAChR) recognition by RABV (Lentz, 1990). RABV-G residues Asn194, Arg199, Gln244, and Thr245 form extensive hydrogen bond networks with 1112-1 CDR residues including Asn52, Asn55, Ser101, Asp102, Tyr103, and Asp105. Detailed interactions are provided in Supplementary Fig. 9. **(C)** 17C7 has been observed to bind mostly to the CD in addition to a small contact with the PHD. Notably, 17C7 engages residues Asn336 and Arg346. Mutations at these locations have been shown to confer resistance to 17C7 neutralization (Sloan et al., 2007; Wang et al., 2011). The 17C7 light chain also forms hydrogen bonds with Lys330 and Arg333, which have been reported to play roles in recognition of the receptor p75NTR (Tuffereau et al., 1998) and in neuroinvasion by rabies virus (Coulon et al., 1998). Detailed interactions are provided in Supplementary Fig. 10. **(D) –(E) Locking of RABV-G in pre-fusion conformation by 17C7 and RVC58**. SPR traces demonstrating that, when RABV-G is captured by site-III-binding mAbs 17C7 or RVC58 respectively at pH 7.4, followed by incubation at pH 5.6, binding of the pre-fusion conformation-specific RVC20 Fab remains possible. RVC20 and RVC58 are specific for the pre-fusion conformation: RVC58 fails to capture RABV-G at pH 5.6, and RVC20 does not bind at pH 5.6 unless RABV-G had previously been captured by 17C7 in pre-fusion conformation.

The mAb 1112-1 epitope encompasses three β-strands and five loops in the PHD (Fig. 4B), described in detail in Supplementary Fig. 8–9. The epitope includes but also extends well beyond the classically-described site II (residues 34–42 and 198–200) (Kuzmina et al., 2013), and overlaps with that of mAb RVC20 (as visualized in a previous crystal structure of RVC20 in complex with the isolated RABV-G PHD) (Fig. 4A and Supplementary Fig. 6) (Hellert et al., 2020). We confirmed by competition ELISA that 1112-1 and RVC20 cannot bind RABV-G simultaneously (Supplementary Fig. 6C).

The mAb 17C7 epitope encompasses both the CD and to a lesser extent the PHD (Fig. 4C, Supplementary Fig. 8 and 10). It includes Asn336 and Arg346 in RABV-G, mutation of which have been shown to confer relative resistance to 17C7 neutralization in the CVS strain of the virus (Sloan et al., 2007; Wang et al., 2011): our structure reveals that these residues form hydrogen bonds with 17C7 paratope residues. The epitope also includes several additional residues which vary between phylogroup I lyssavirus species residues (Supplementary Fig. 8). The mAb 17C7 light chain also forms hydrogen bonds with Lys330 and Arg333, which have been reported to be important for recognition of the neurotrophin receptor, p75NTR, and in neuroinvasion by rabies virus (Coulon et al., 1998; Tuffereau et al., 1998).

Given the clinical importance of 17C7, we sought to explore its mechanism of action. Due to the presence of Arg333 within site III, and the known importance of this residue in neurovirulence and p75NTR binding, it was previously postulated that site-III-binding antibodies may function by blocking RABV-G binding to p75NTR (Dietzschold et al., 1983; Yang et al., 2020). However, p75NTR is dispensable for RABV-G pathogenicity (Tuffereau et al., 2007), suggesting this is unlikely to be the only mechanism of action of such antibodies. In contrast, the site I-binding mAb RVC20 has previously been suggested to act in a receptor-independent manner, by ‘locking’ RABV-G in pre-fusion conformation. While RVC20 bound (post-fusion) RABV-G poorly at low pH, the rate of RVC20 dissociation from RABV-G was pH-independent if association had occurred at neutral pH (Hellert et al., 2020).

Mapping the 17C7 epitope onto the low-pH-derived G^ecto^ structure (Yang et al., 2020) revealed that, although the contact residues are likely to remain largely accessible in the post-fusion conformation, the acidic-pH-induced structural transition may separate the major (CD) and minor (PHD) regions of the bipartite epitope by more than 20 Å (Supplementary Fig. 6B). We hypothesized that binding of 17C7 may hinder this separation and hence prevent the pre-to post-fusion conformational transition in RABV-G.

We confirmed the conformational specificity of RVC20, 1112-1, and 17C7 (site I, II and III mAbs respectively), using a surface plasmon resonance (SPR) assay (Supplementary Fig. 11). Consistent with our and other data (Hellert et al., 2020), RVC20 binding was abrogated at low pH. Despite appreciable reduction in their binding affinity as compared to that at pH 7.4, 1112-1 and 17C7 remained capable of binding RABV-G with nanomolar affinity at pH 5.6 (Supplementary Fig. 11D). In the case of 17C7, this is consistent with the fact that both CD and PHD elements of the binding site remain accessible on the RABV-G^ecto^ structure, and PHD makes a relatively minor contribution to the binding footprint.

As 17C7 binding in itself could not be used as a reporter of RABV-G conformation, we used a ‘sandwich-configuration’ SPR assay to assess the effect of 17C7, or another site-III binding mAb (RVC58 (De Benedictis et al., 2016)) upon pH-induced RABV-G conformational change (Fig. 4D–E). When RABV-G had been captured by either 17C7 or RVC58 under neutral conditions, followed by lowering of the pH to 5.6 for 300s, RVC20 remained able to bind despite being applied at pH 5.6. The conformational change which abrogates RVC20 binding thus cannot take place when RABV-G has already been bound by 17C7 or RVC58 in pH-neutral / pre-fusion conformation. Therefore, although binding of 17C7 is possible in both pre- and post-fusion conformations of RABV-G, the presence of 17C7 blocks the transition between conformations.

## Discussion

Given the health and economic burden of rabies disease and recent progression in vaccine technology, there is both need and opportunity to develop more immunogenic and more cost-effective rabies vaccines. Design of highly-expressed and stable pre-fusion RABV-G trimers would considerably assist production of both recombinant protein and nucleic-acid- or viral-vectored vaccines. The structure we present here resolves the RABV-G trimerization interface, L4 linker and likely authentic pre-fusion conformation of L5. Each of these elements plays an important role in maintenance of the pre-fusion trimeric architecture and transition to post-fusion conformation, and is likely to be a target of structure-guided immunogen design. Our findings that H261A/L and H270P substitutions increase pre-fusion conformational stability are important initial steps towards this goal.

We also present structures of protective antibodies bound to two of RABV-G’s three major antigenic sites. Notably, this new data makes clear the structural basis of the known vulnerability of the licensed therapeutic, 17C7, to polymorphisms between RABV isolates and between RABV and other lyssaviruses (including bat lyssaviruses within the relatively-closely-related phylogroup I, which are known to cause human disease) (De Benedictis et al., 2016). This should facilitate the development of future more broadly nAbs, and highlights the risks of reliance for protection on a single antibody, even against a relatively conserved antigen such as RABV-G.

Finally, we show that two site-III-binding nAbs (17C7 and RVC58) are capable of locking RABV-G in the pre-fusion conformation. In contrast with the previous suggestion that site-III-binding antibodies may act by blockade of RABV-G — p75NTR interaction, which is dispensable for virulence (Hellert et al., 2020; Yang et al., 2020), antibody-mediated blockade of the conformational transition is likely to be effective against entry to all cell types. This builds upon similar previous findings with the site-I-binding antibody RVC20, and raises the intriguing question of whether blockade of conformational transition may be a mechanism shared by many, if not all, RABV-G-binding nAbs. Broader understanding of the mechanism of action of this important therapeutic class would be assisted by further structural and functional data regarding RABV-G’s interactions with the multiple proteins which may act as host receptors.

Together, our findings will facilitate rational development of improved vaccine immunogens and therapeutics against this major human and animal pathogen, and extend understanding of the mechanism by which protective antibodies can neutralize the virus.

## Acknowledgments

We are grateful for support and advice from Matthew Higgins and Adam Ritchie. The 1112-1 hybridoma was a kind gift of Hildegund Ertl, Wistar Institute. The pVIP-ENTR plasmid was a kind gift of Dr Martino Bardelli, Jenner Institute. We acknowledge access to cryo-EM facilities at the U.K. National Electron Bio-Imaging Centre (eBIC) at the Diamond Light Source, funded by the Wellcome Trust, MRC, and BBSRC. We are grateful to Vinod Vogirala, Helen Duyvesteyn, and Loic Carrique for assistance in data acquisition at the eBIC and the Oxford Particle Imaging Centre.

## Author contributions

Conceptualization: ADD, SF, TAB, GA

Methodology: ADD, SF, TAB, WMN

Investigation: All

Visualization: ADD, SF, TAB, WMN

Funding acquisition: ADD, SF

Project administration: ADD

Supervision: ADD

Writing–original draft: ADD, TAB, WMN, SF

Writing–review & editing: All

## Data and materials availability

All data needed to evaluate the conclusions in the paper are present in the paper and/or the Supplementary Materials.

Cryo-EM density map with the corresponding atomic coordinates for RABV-G – Fab 17C7 – Fab 1112-1 complex will be deposited in the Electron Microscopy Data Bank and Protein Data Bank with the accession codes EMD-XXXXX and YYYY, upon initial revision of the manuscript.

## References

(2018). WHO Expert Consultation on Rabies, Third Report, Vol 1012 (Geneva: World Health Organization).

Albertini, A.A., Baquero, E., Ferlin, A., and Gaudin, Y. (2012). Molecular and cellular aspects of rhabdovirus entry. Viruses 4, 117–139.

Aldrich, C., Leroux-Roels, I., Huang, K.B., Bica, M.A., Loeliger, E., Schoenborn-Kellenberger, O., Walz, L., Leroux-Roels, G., von Sonnenburg, F., and Oostvogels, L. (2021). Proof-of-concept of a low-dose unmodified mRNA-based rabies vaccine formulated with lipid nanoparticles in human volunteers: A phase 1 trial. Vaccine 39, 1310–1318.

Baquero, E., Albertini, A.A., Raux, H., Buonocore, L., Rose, J.K., Bressanelli, S., and Gaudin, Y. (2015). Structure of the low pH conformation of Chandipura virus G reveals important features in the evolution of the vesiculovirus glycoprotein. PLoS Pathog 11, e1004756.

Baquero, E., Buonocore, L., Rose, J.K., Bressanelli, S., Gaudin, Y., and Albertini, A.A. (2012). Crystallization and preliminary X-ray analysis of Chandipura virus glycoprotein G. Acta Crystallogr Sect F Struct Biol Cryst Commun 68, 1094–1097.

Benmansour, A., Leblois, H., Coulon, P., Tuffereau, C., Gaudin, Y., Flamand, A., and Lafay, F. (1991). Antigenicity of rabies virus glycoprotein. J Virol 65, 4198–4203.

Coulon, P., Ternaux, J.P., Flamand, A., and Tuffereau, C. (1998). An avirulent mutant of rabies virus is unable to infect motoneurons in vivo and in vitro. J Virol 72, 273–278.

De Benedictis, P., Minola, A., Rota Nodari, E., Aiello, R., Zecchin, B., Salomoni, A., Foglierini, M., Agatic, G., Vanzetta, F., Lavenir, R., et al. (2016). Development of broad-spectrum human monoclonal antibodies for rabies post-exposure prophylaxis. EMBO Mol Med 8, 407–421.

Dietzschold, B., Kao, M., Zheng, Y.M., Chen, Z.Y., Maul, G., Fu, Z.F., Rupprecht, C.E., and Koprowski, H. (1992). Delineation of putative mechanisms involved in antibody-mediated clearance of rabies virus from the central nervous system. Proc Natl Acad Sci U S A 89, 7252–7256.

Dietzschold, B., Wunner, W.H., Wiktor, T.J., Lopes, A.D., Lafon, M., Smith, C.L., and Koprowski, H. (1983). Characterization of an antigenic determinant of the glycoprotein that correlates with pathogenicity of rabies virus. Proc Natl Acad Sci U S A 80, 70–74.

Ertl, H.C.J. (2019). New Rabies Vaccines for Use in Humans. Vaccines (Basel) 7.

FDA (2021). Rabies: Developing Monoclonal Antibody Cocktails for the Passive Immunization Component of Post-Exposure Prophylaxis Guidance for Industry (Draft).

Ferlin, A., Raux, H., Baquero, E., Lepault, J., and Gaudin, Y. (2014). Characterization of pH-sensitive molecular switches that trigger the structural transition of vesicular stomatitis virus glycoprotein from the postfusion state toward the prefusion state. J Virol 88, 13396–13409.

Fooks, A.R., Cliquet, F., Finke, S., Freuling, C., Hemachudha, T., Mani, R.S., Müller, T., Nadin-Davis, S., Picard-Meyer, E., Wilde, H., et al. (2017). Rabies. Nature Reviews Disease Primers 3, 17091.

Gaudin, Y., Ruigrok, R.W., Tuffereau, C., Knossow, M., and Flamand, A. (1992). Rabies virus glycoprotein is a trimer. Virology 187, 627–632.

Gaudin, Y., Tuffereau, C., Durrer, P., Brunner, J., Flamand, A., and Ruigrok, R. (1999). Rabies virus-induced membrane fusion. Mol Membr Biol 16, 21–31.

Gaudin, Y., Tuffereau, C., Segretain, D., Knossow, M., and Flamand, A. (1991). Reversible conformational changes and fusion activity of rabies virus glycoprotein. J Virol 65, 4853–4859.

Gogtay, N.J., Munshi, R., Ashwath Narayana, D.H., Mahendra, B.J., Kshirsagar, V., Gunale, B., Moore, S., Cheslock, P., Thaker, S., Deshpande, S., et al. (2018). Comparison of a Novel Human Rabies Monoclonal Antibody to Human Rabies Immunoglobulin for Postexposure Prophylaxis: A Phase 2/3, Randomized, Single-Blind, Noninferiority, Controlled Study. Clinical infectious diseases : an official publication of the Infectious Diseases Society of America 66, 387–395.

Hampson, K., Coudeville, L., Lembo, T., Sambo, M., Kieffer, A., Attlan, M., Barrat, J., Blanton, J.D., Briggs, D.J., Cleaveland, S., et al. (2015). Estimating the global burden of endemic canine rabies. PLoS Negl Trop Dis 9, e0003709.

Hellert, J., Buchrieser, J., Larrous, F., Minola, A., de Melo, G.D., Soriaga, L., England, P., Haouz, A., Telenti, A., Schwartz, O., et al. (2020). Structure of the prefusion-locking broadly neutralizing antibody RVC20 bound to the rabies virus glycoprotein. Nat Commun 11, 596.

Koraka, P., Bosch, B.J., Cox, M., Chubet, R., Amerongen, G., Lovgren-Bengtsson, K., Martina, B.E., Roose, J., Rottier, P.J., and Osterhaus, A.D. (2014). A recombinant rabies vaccine expressing the trimeric form of the glycoprotein confers enhanced immunogenicity and protection in outbred mice. Vaccine 32, 4644–4650.

Kuzmina, N.A., Kuzmin, I.V., Ellison, J.A., and Rupprecht, C.E. (2013). Conservation of Binding Epitopes for Monoclonal Antibodies on the Rabies Virus Glycoprotein. Journal of Antivirals & Antiretrovirals 5, 37–43.

Lentz, T.L. (1990). Rabies virus binding to an acetylcholine receptor alpha-subunit peptide. Journal of molecular recognition : JMR 3, 82–88.

Lentz, T.L., Burrage, T.G., Smith, A.L., Crick, J., and Tignor, G.H. (1982). Is the acetylcholine receptor a rabies virus receptor? Science 215, 182–184.

Muller, T., Dietzschold, B., Ertl, H., Fooks, A.R., Freuling, C., Fehlner-Gardiner, C., Kliemt, J., Meslin, F.X., Franka, R., Rupprecht, C.E., et al. (2009). Development of a mouse monoclonal antibody cocktail for post-exposure rabies prophylaxis in humans. PLoS Negl Trop Dis 3, e542.

Ren, J., Wen, L., Gao, X., Jin, C., Xue, Y., and Yao, X. (2009). DOG 1.0: illustrator of protein domain structures. Cell Res 19, 271–273.

Roche, S., Bressanelli, S., Rey, F.A., and Gaudin, Y. (2006). Crystal structure of the low-pH form of the vesicular stomatitis virus glycoprotein G. Science 313, 187–191.

Roche, S., Rey, F.A., Gaudin, Y., and Bressanelli, S. (2007). Structure of the prefusion form of the vesicular stomatitis virus glycoprotein G. Science 315, 843–848.

Si, Z., Zhang, J., Shivakoti, S., Atanasov, I., Tao, C.L., Hui, W.H., Zhou, K., Yu, X., Li, W., Luo, M., et al. (2018). Different functional states of fusion protein gB revealed on human cytomegalovirus by cryo electron tomography with Volta phase plate. PLoS Pathog 14, e1007452.

Sloan, S.E., Hanlon, C., Weldon, W., Niezgoda, M., Blanton, J., Self, J., Rowley, K.J., Mandell, R.B., Babcock, G.J., Thomas, W.D., Jr., et al. (2007). Identification and characterization of a human monoclonal antibody that potently neutralizes a broad panel of rabies virus isolates. Vaccine 25, 2800–2810.

Sparrow, E., Torvaldsen, S., Newall, A.T., Wood, J.G., Sheikh, M., Kieny, M.P., and Abela-Ridder, B. (2019). Recent advances in the development of monoclonal antibodies for rabies post exposure prophylaxis: A review of the current status of the clinical development pipeline. Vaccine 37 Suppl 1, A132–A139.

Thoulouze, M.I., Lafage, M., Schachner, M., Hartmann, U., Cremer, H., and Lafon, M. (1998). The neural cell adhesion molecule is a receptor for rabies virus. J Virol 72, 7181–7190.

Tuffereau, C., Benejean, J., Blondel, D., Kieffer, B., and Flamand, A. (1998). Low-affinity nerve-growth factor receptor (P75NTR) can serve as a receptor for rabies virus. EMBO J 17, 7250–7259.

Tuffereau, C., Schmidt, K., Langevin, C., Lafay, F., Dechant, G., and Koltzenburg, M. (2007). The rabies virus glycoprotein receptor p75NTR is not essential for rabies virus infection. J Virol 81, 13622–13630.

Wang, J., Wang, Z., Liu, R., Shuai, L., Wang, X., Luo, J., Wang, C., Chen, W., Wang, X., Ge, J., et al. (2018). Metabotropic glutamate receptor subtype 2 is a cellular receptor for rabies virus. PLoS Pathog 14, e1007189.

Wang, Y., Rowley, K.J., Booth, B.J., Sloan, S.E., Ambrosino, D.M., and Babcock, G.J. (2011). G glycoprotein amino acid residues required for human monoclonal antibody RAB1 neutralization are conserved in rabies virus street isolates. Antiviral Res 91, 187–194.

Wiktor, T.J., Gyorgy, E., Schlumberger, D., Sokol, F., and Koprowski, H. (1973). Antigenic properties of rabies virus components. J Immunol 110, 269–276.

Yamada, K., Noguchi, K., Nonaka, D., Morita, M., Yasuda, A., Kawazato, H., and Nishizono, A. (2013). Addition of a single N-glycan to street rabies virus glycoprotein enhances virus production. J Gen Virol 94, 270–275.

Yang, F., Lin, S., Ye, F., Yang, J., Qi, J., Chen, Z., Lin, X., Wang, J., Yue, D., Cheng, Y., et al. (2020). Structural Analysis of Rabies Virus Glycoprotein Reveals pH-Dependent Conformational Changes and Interactions with a Neutralizing Antibody. Cell host & microbe 27, 441–453 e447.

