## Supplementary Figures and Tables for "Structure of trimeric pre-fusion rabies virus glycoprotein in complex with two protective antibodies"

##### Figure S1: Cryo-EM processing flowchart of RABV-G in complex with Fabs 17C7 and 1112-1

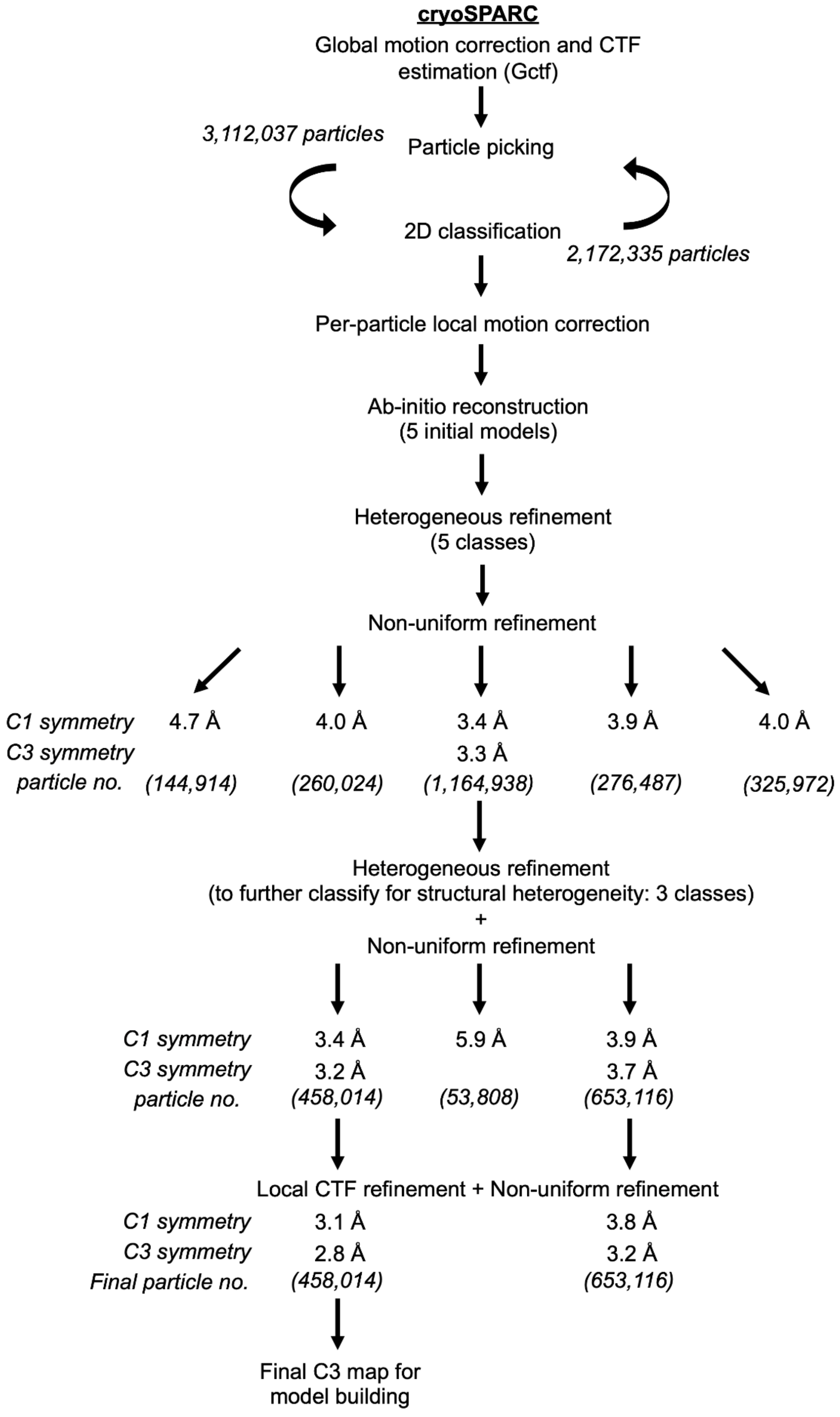


**Fig. S1. Cryo-EM processing flowchart of RABV-G in complex with Fabs 17C7 and 1112-1**. All processing steps were performed in cryoSPARC (Punjani et al., 2017).

#### Figure S2: Resolution analysis of the cryo-EM structure of RABV-G in complex with Fabs 17C7 and 1112-1

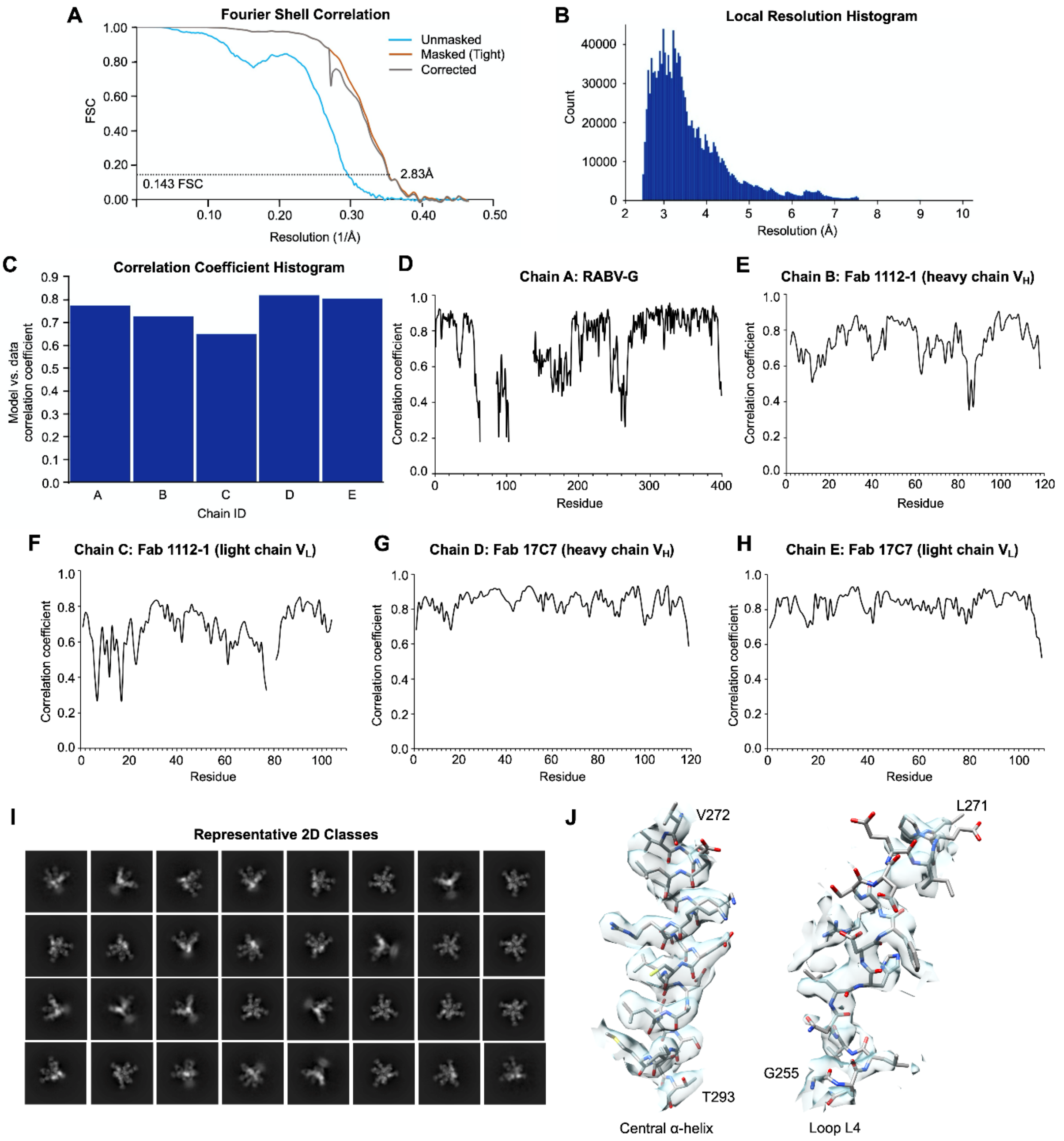


**Fig. S2. Resolution analysis of the cryo-EM structure of RABV-G in complex with Fabs 17C7 and 1112-1.** (**A**) Fourier shell correlation (FSC) plots of reconstructions using gold-standard refinement in cryoSPARC (Punjani et al., 2017). Map resolutions were determined according to the 0.143 FSC cutoff. Curves are shown for unmasked (cyan), masked (brown) and corrected (grey) maps. (**B**) Local resolution histogram determined by cryoSPARC (Punjani et al., 2017). (**C**) Correlation coefficient per chain histogram determined by Phenix (Adams et al., 2002). (**D**–**H**) Correlation coefficient per residue graphs for (**D**) RABV-G, (**E**) Fab 1112-1 heavy chain $V_H$, (**F**) Fab 1112-1 light chain $V_L$, (**G**) Fab 17C7 heavy chain $V_H$, (**H**) Fab 17C7 light chain $V_L$. (**I**) Representative 2D classes used in 3D reconstruction. (**J**) Representative fit of atomic model into density.

#### Figure S3: Contacts of the L5 linker with CD and FD

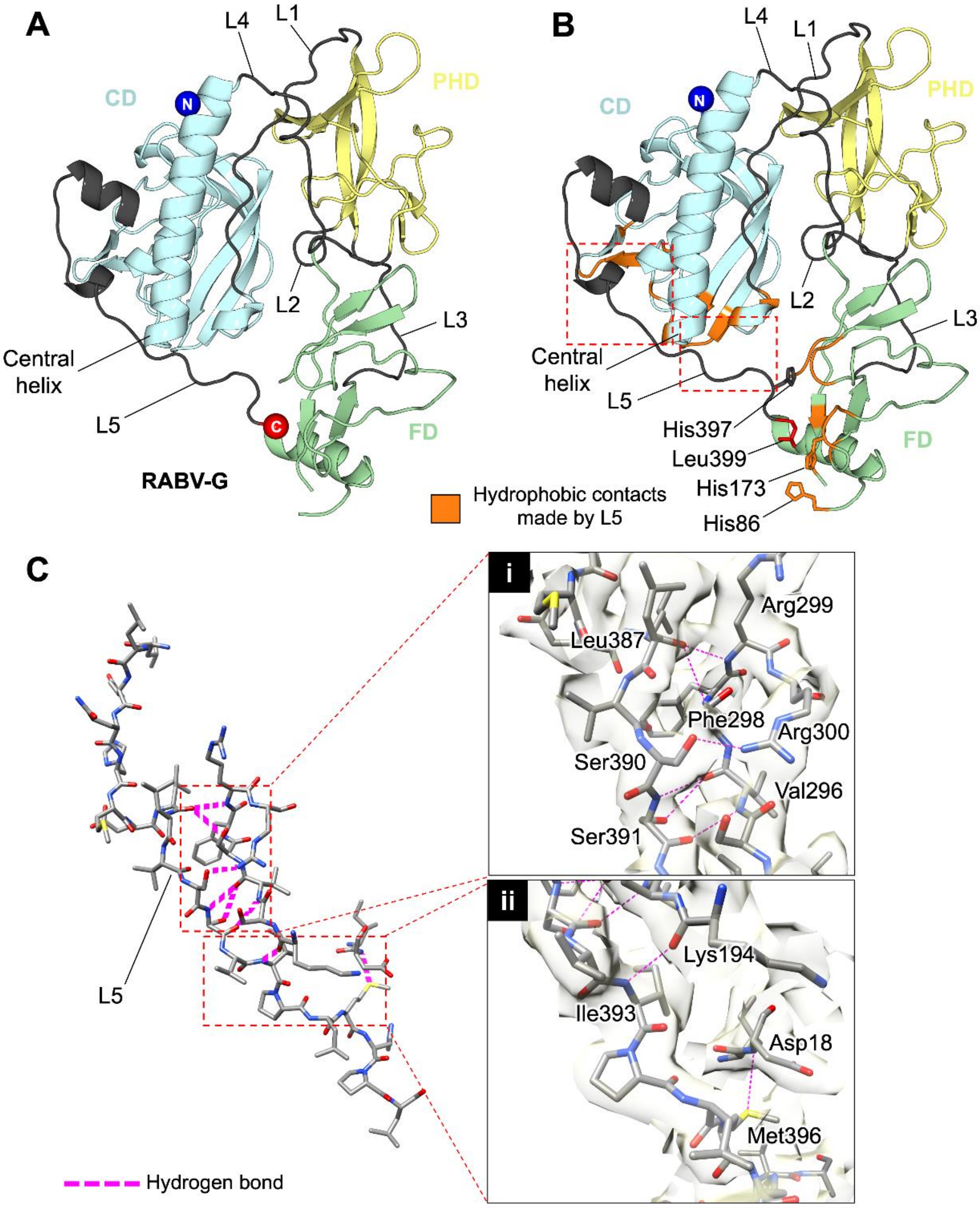


**Fig. S3. Contacts of the L5 linker with CD and FD.**

(**A**) Structure of a protomer of the RABV-G ectodomain crown is displayed in a cartoon representation, with PHD, CD, and FD colored yellow, cyan, and green, respectively. Inter-domain linkers are colored dark gray. The N and C-termini of the structure are shown as spheres and colored blue and red, respectively.

(**B**) Structure of RABV-G protomer is shown as in panel A. Residues involved in intra-protomeric hydrophobic interactions with L5 are colored orange. Dashed line boxes indicate regions where L5 forms intra-protomeric hydrogen bonds. These regions are detailed in panel C.

(**C**) Intra-protomeric hydrogen bonds mediated by L5. Residues are shown in stick representation with carbon, nitrogen, oxygen, and sulfur atoms colored gray, blue, red, and yellow, respectively. Detailed interactions as indicated by the dashed line boxes are enlarged in sub-panels i–ii. Residues forming hydrogen bonds (pink dashed lines) are labeled. The cryo-EM map is shown with partial transparency. Interactions were determined using ePISA (Krissinel and Henrick, 2007).

Figure S4: L4 intra-protomeric interactions

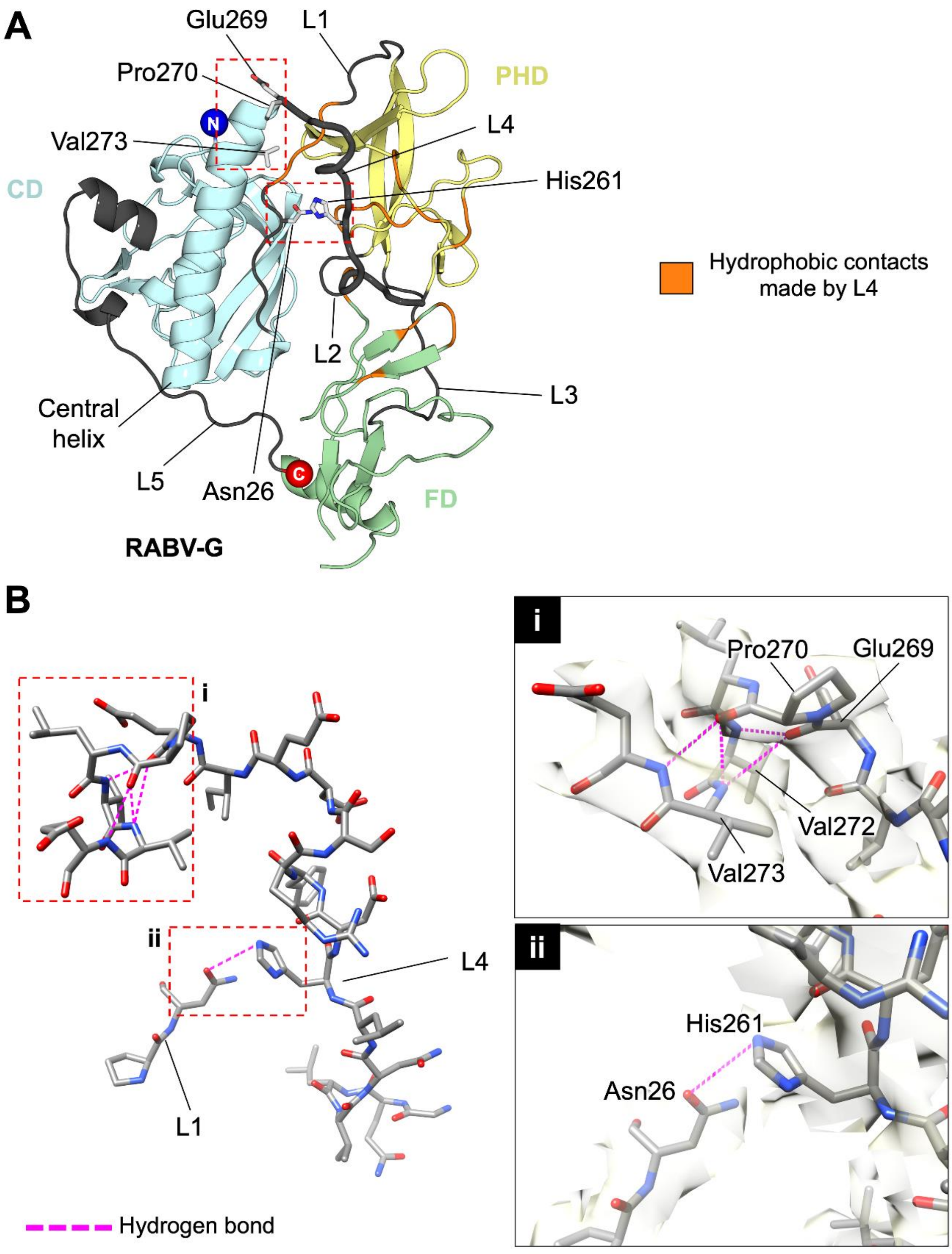


**Fig. S4. Linker L4 intra-protomeric contacts.** (**A**) Structure of the RABV-G crown ectodomain region monomer is shown as in *Fig. S3, panel A*. L4 is labeled and shown with thicker width. Residues involved in intra-protomeric hydrophobic interactions with L4 are colored orange while residues involved in hydrogen bonding with L4 are shown as sticks. (**B**) Detailed visualization of intra-protomeric hydrogen bonds formed by L4. Residues are displayed as sticks with carbon, nitrogen, and oxygen atoms colored gray, blue, and red, respectively. Detailed interactions as indicated by the dashed-line boxes are enlarged in sub-panels i–ii. Residues forming hydrogen bonds (pink dashed lines) are labeled. The cryo-EM map is shown with partial transparency. Interactions were determined using ePISA (Krissinel and Henrick, 2007).

#### Figure S5: RABV-G structure alignment with VSV-G crystal structure

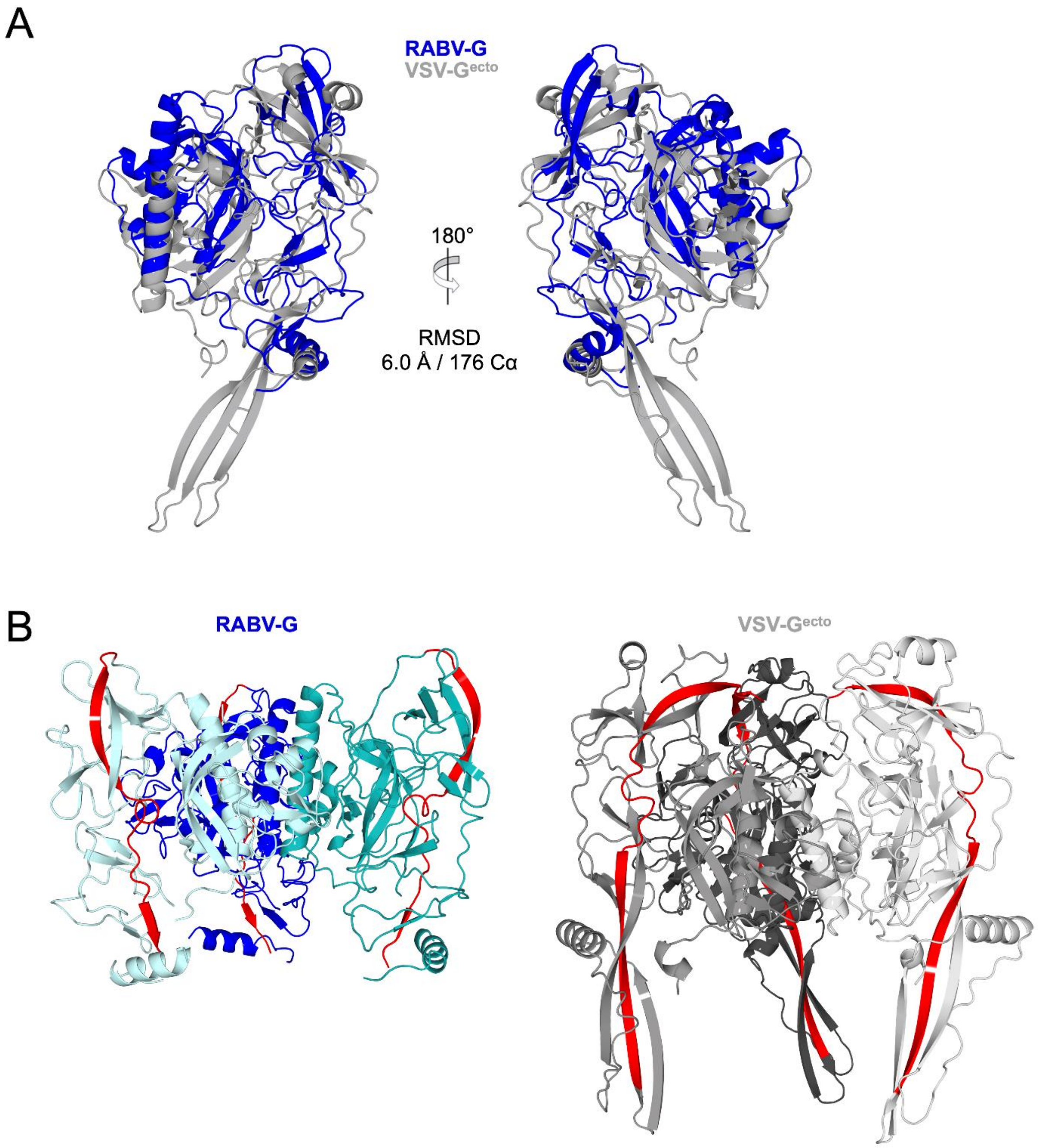


**Fig. S5. Structural comparison of RABV-G and VSV-G.**

(**A**) Structural superimposition of a single RABV-G protomer with VSV-G pre-fusion ectodomain protomer (PDB ID: 5I2S) (Roche et al., 2007). Structures are displayed in cartoon representation with RABV-G colored blue and VSV-G colored gray. Root-mean-square deviation (RMSD) of the alignment is indicated. Despite the relatively high RMSD of 6.0 Å, the overall conservation of the domain architecture can be seen.

(**B**) Differential angulations of RABV-G and VSV-G in the context of a trimer assembly. Trimeric RABV-G and VSV-$G^{ecto}$ are displayed as blue and gray cartoons, with different color shade for each protomer. Residues 36–63 in RABV-G and residues 36–69 in VSV-$G^{ecto}$ are colored red to show the angulations of the G molecules.

#### Figure S6: Antigenic sites plotted on surface of RABV-G

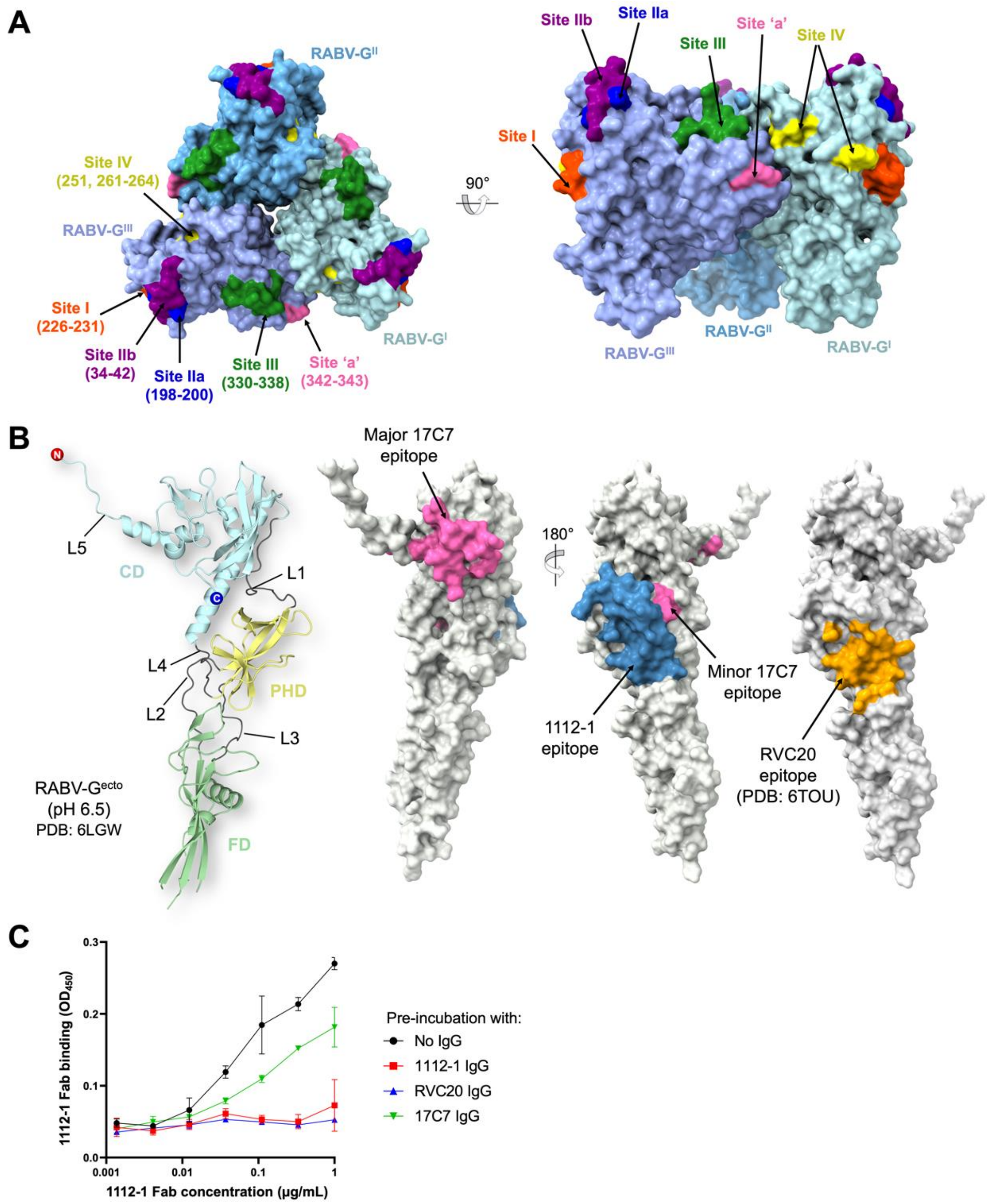


**Fig. S6. RABV-G antigenic sites.**

(**A**) Characterized antigenic sites I–IV and 'a' (Kuzmina et al., 2013) are mapped onto the trimeric RABV-G structure (surface representation, with each protomer in different shades of blue). Residue numbers for each site are indicated in parentheses.

(**B**) (*Left*) Crystal structure of RABV-G$^{ecto}$ obtained at pH 6.5 and hence believed to represent post-fusion conformation (PDB ID: 6LGW) (Yang et al., 2020) displayed as cartoon with CD, PHD, FD, and L1–L5 colored and labeled accordingly. (*Right*) Footprints of 17C7 and 1112-1 epitopes determined in this study are mapped onto the RABV-G$^{ecto}$ (pH 6.5) crystal structure (white, surface representation). 17C7 footprint, pink; 1112-1 footprint, blue. Residues previously shown to form contacts with RVC20 are shown in orange – this epitope is not believed to be accessible in post-fusion conformation (Hellert et al., 2020).

**(C)** Binding of 1112-1 IgG to pre-fusion RABV-G is compatible with binding of 17C7, but not RVC20, as demonstrated by ELISA. Plates coated with recombinant RABV-G–C-tag were pre-incubated with 20 μg/mL of the indicated antibodies, before application of TwinStrep-tagged 1112-1 Fab and detection with horseradish-peroxidase-conjugated Streptactin. Points and error bars represent median and range of triplicate wells (technical replicates).

#### Figure S7: Densities corresponding to asparagine-linked glycosylation sequons

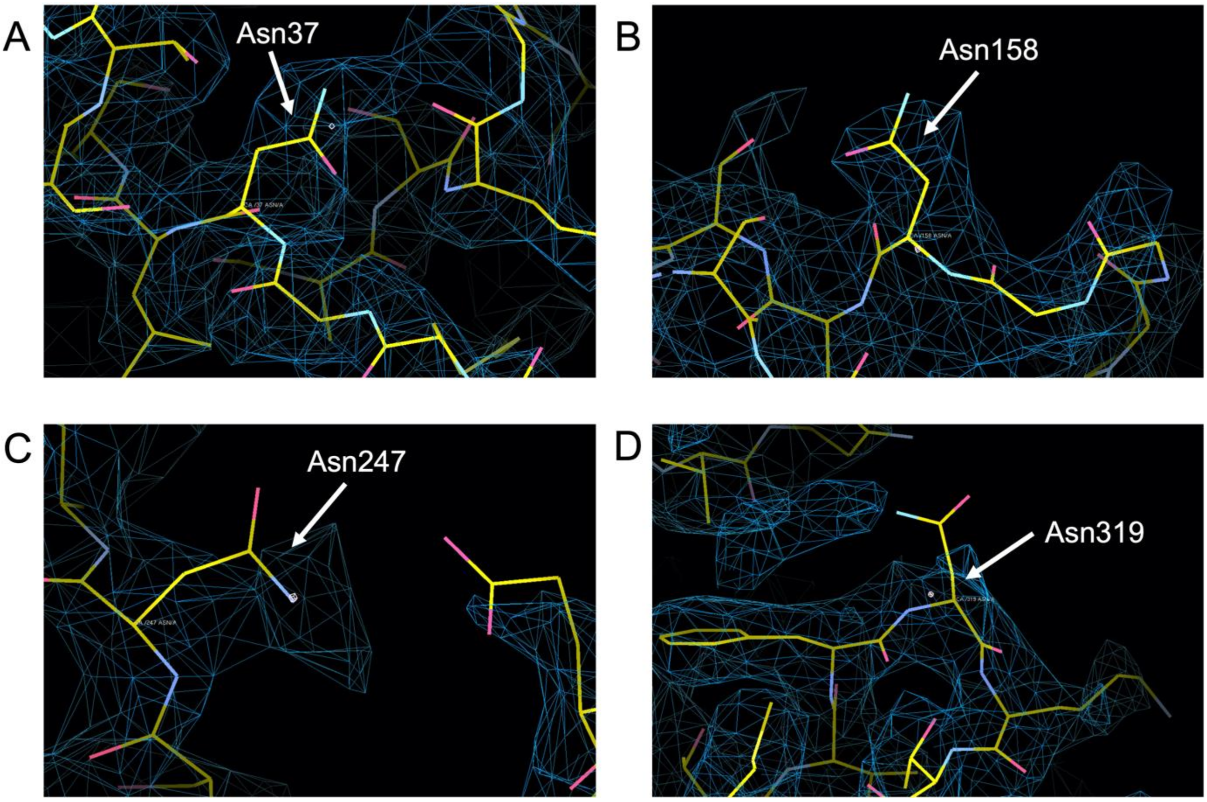


**Fig. S7. Densities corresponding to asparagine (N)-linked glycosylation sequons within RABV-G.**

(**A**)–(**D**) Atomic model of RABV-G displaying N-linked glycosylation sequons (NXT/S, X ≠ P). Amino acid residues are shown in stick representation, with carbon, nitrogen, oxygen, and sulfur atoms colored yellow, blue, pink, and dark yellow, respectively. Asn37, Asn158, Asn247, and Asn319 are labeled. Map densities are shown as blue mesh, rendered in Coot (Emsley and Cowtan, 2004). No ordered densities corresponding to glycans were observed in our map, but we are unable to distinguish between disordered glycans and unoccupied sequons.

#### Figure S8: Conservation and variation of 17C7 and 1112-1 contact residues across lyssaviruses

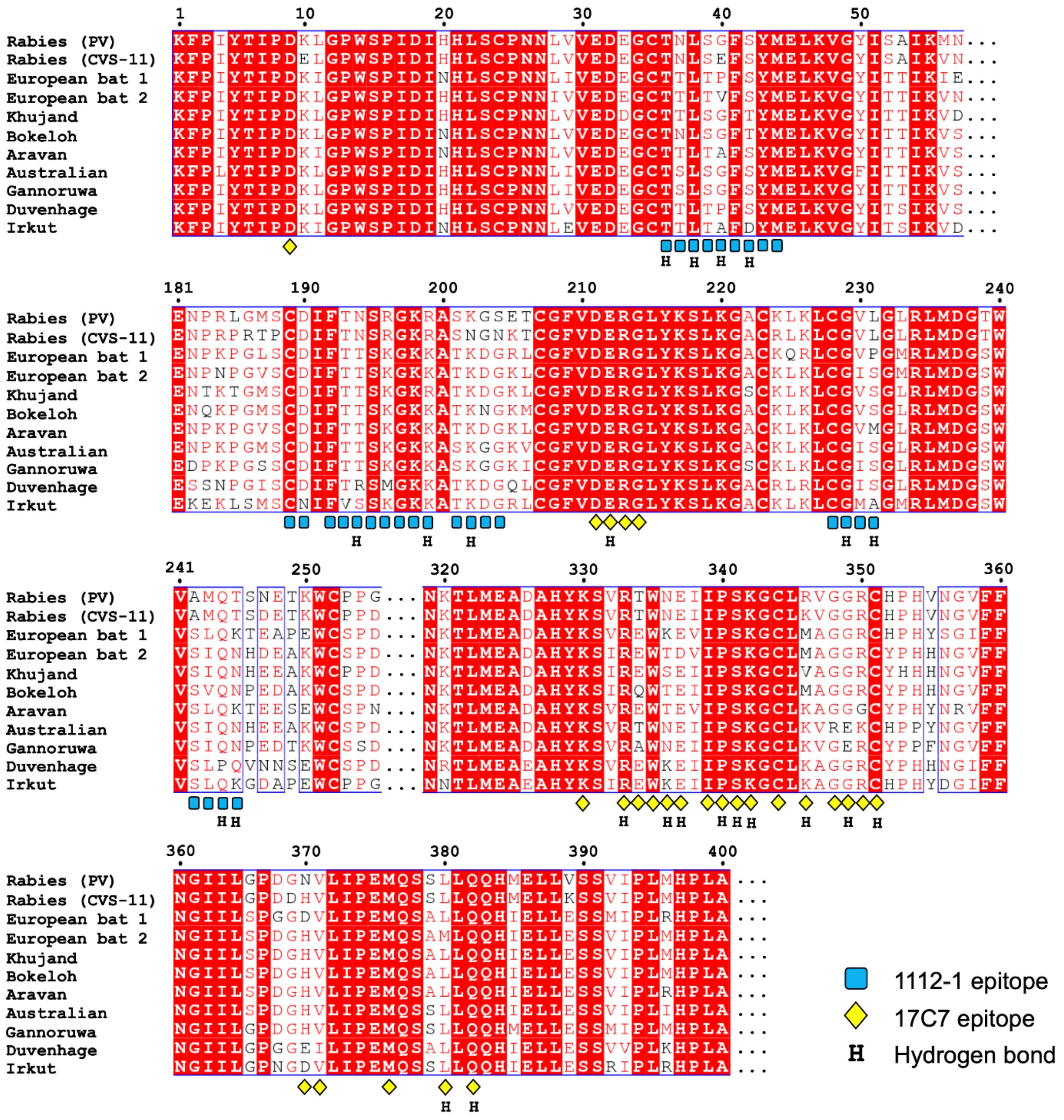


**Fig. S8. Conservation and variation of 17C7 and 1112-1 contact residues across lyssaviruses.** Glycoprotein amino acid sequences of PV strain rabies virus(Kim et al., 2016), CVS-11 rabies virus (ADJ29911.1), European bat lyssavirus 1 (YP_001285391.1), European bat lyssavirus 2 (YP_001285396.1), Khujand virus (YP_009094330.1), Bokeloh bat lyssavirus (YP_009091812.1), Aravan lyssavirus (YP_007641395.1), Australian bat lyssavirus (QIN55368.1), Gannoruwa lyssavirus (YP_009325517.1), Duvenhage lyssavirus (YP_007641405.1), and Irkut lyssavirus (AFP74571.1) were determined using MultAlin(Corpet, 1988) and plotted with ESPript(Gouet et al., 2003). Identical residues are shaded in red. Residues interacting with Fabs 1112-1 and 17C7 in our structure were identified using ePISA (Krissinel and Henrick, 2007) and are annotated beneath the alignment as indicated. Residues forming hydrogen bonds with the Fab fragments are denoted 'H'.

#### Figure S9: Representation of RABV-G – 1112-1 contacts

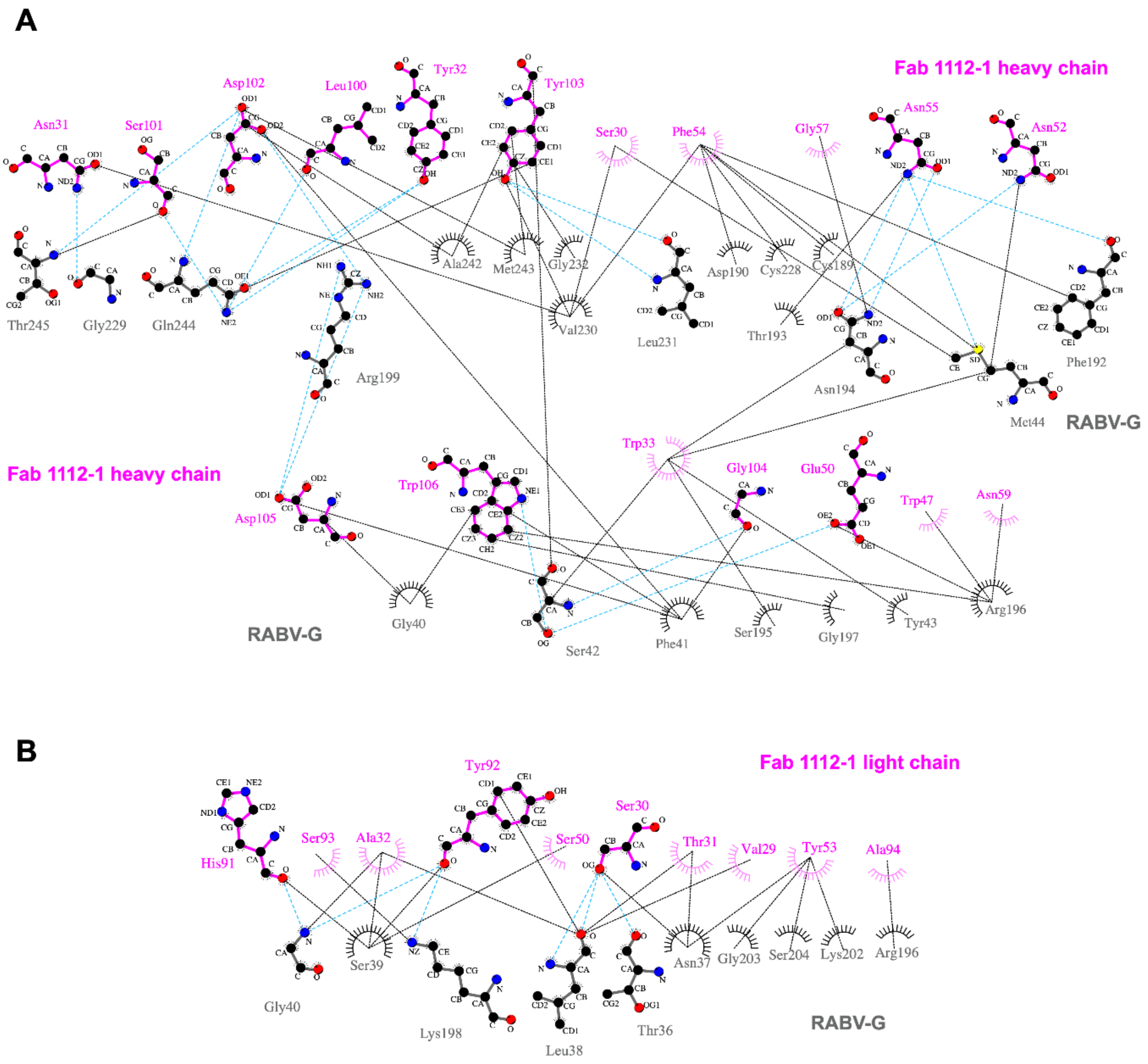


**Fig. S9. Schematic diagram of contacts formed at the RABV-G – Fab 1112-1 interface.** (**A**) RABV-G – Fab 1112-1 heavy chain CDR interactions. (*Lower*) RABV-G–1112-1 light chain CDR interactions. RABV-G residues are colored gray and Fab 1112-1 residues are colored magenta. Atoms corresponding to carbon, nitrogen, and oxygen are shown as black, blue, and red balls, respectively. Residues involved in hydrogen bonding are shown as sticks; residues involved in hydrophobic interactions are shown as spoked arcs. Hydrogen bonds and hydrophobic interactions are shown as cyan and black dotted lines, respectively. Plots were generated with LigPlot+ (Laskowski and Swindells, 2011).

Figure S10: Representation of RABV-G – 17C7 contacts

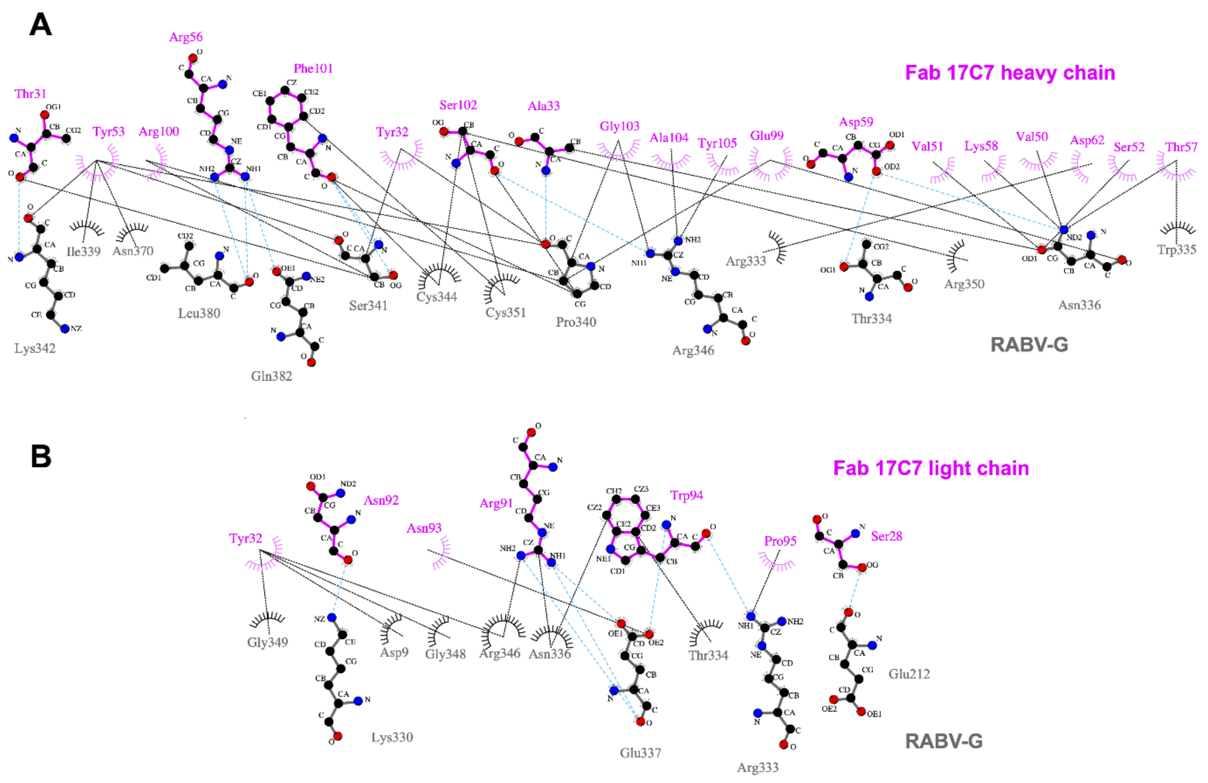


**Fig. S10. Schematic diagram of contacts formed at the RABV-G – Fab 17C7 interface.** (**A**) RABV-G–17C7 heavy chain interactions. (*Lower*) RABV-G–17C7 light chain interactions. RABV-G residues are colored gray and Fab 17C7 residues are colored magenta. Atoms corresponding to carbon, nitrogen, and oxygen are shown as black, blue, and red balls, respectively. Residues involved in hydrogen bonding are shown as sticks; residues involved in hydrophobic interactions are shown as spoked arcs. Hydrogen bonds and hydrophobic interactions are shown as cyan and black dotted lines, respectively. Plots were generated with LigPlot+ (Laskowski and Swindells, 2011).

#### Figure S11: Kinetics of RABV-G interactions with Fabs under acidic and neutral pH conditions

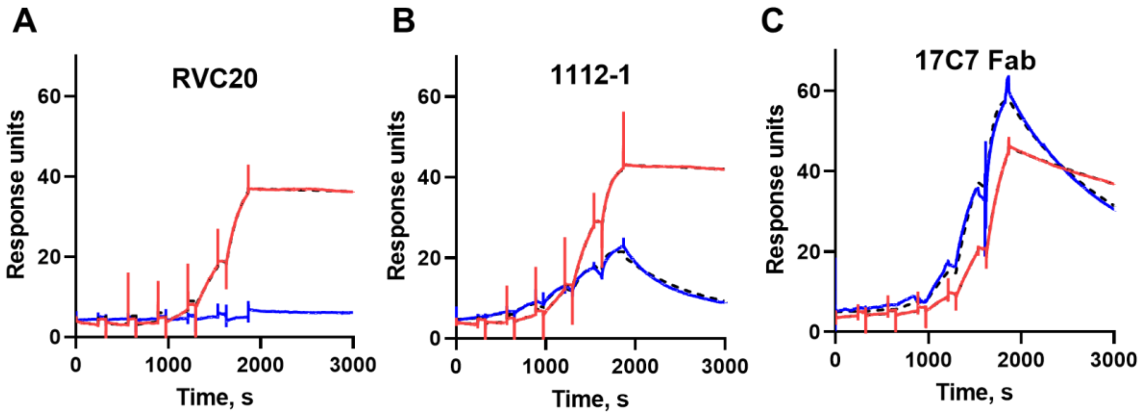


**D**

| Fab | Buffer pH | ka (1/Ms) | kd (1/s) | $K_D$ (nM) |
|---|---|---|---|---|
| RVC20 | pH 7.5 | 2.0x10$^{5}$ | 2.0x10$^{-5}$ | 0.01 |
| | pH 5.6 | none | none | none |
| 1112-1 | pH 7.5 | 3.8x10$^{5}$ | 2.4x10$^{-5}$ | 0.06 |
| | pH 5.6 | 8.0x10$^{4}$ | 1.2x10$^{-3}$ | 15 |
| 17C7 | pH 7.5 | 1.4x10$^{5}$ | 2.2x10$^{-4}$ | 1.6 |
| | pH 5.6 | 7.8x10$^{4}$ | 9.7x10$^{-4}$ | 13 |

**Fig. S11. Kinetics of RABV-G interactions with Fabs under acidic and neutral pH conditions**

**(A–C)** SPR analysis of the kinetics of RVC20, 1112-1 and 17C7 interactions with RABV-G. In each case data observed at pH 7.4 and pH 5.6 are represented by red and blue lines respectively. Calculated kinetic and affinity values shown in **(D).** Fab concentration ranges used differed between cycles, as described in methods.

### Supplementary Tables

#### Table S1: Cryo-EM data collection, processing, and model refinement statistics

| | |
|---|---|
| **Data collection and processing** | |
| Microscope | Titan Krios |
| Detector | Gatan K3 |
| Voltage (kV) | 300 |
| Recording mode | Super resolution |
| Electron dose ($e^-$/Å$^2$) | 44.4 |
| Defocus range (µm) | –0.8 to –2.6 |
| Frames | 45 |
| Magnification | 81,000 |
| Final map pixel size (Å/px) | 1.06 |
| Symmetry imposed | C3 |
| No. of movies | 12,884 |
| No. of final particles images | 458,014 |
| Map resolution at 0.143 FSC threshold (Å) | 2.8 |
| Map sharpening B-factor (Å$^2$) | –108 |
| **Model refinement** | |
| FSC model vs. map at 0.5 threshold (Å) | 3.0 |
| CC model vs. map (masked) | 0.80 |
| *Model composition* | |
| Non-hydrogen atoms | 6,130 |
| Protein residues | 790 |
| Non-protein residues | 0 |
| *R.m.s. deviations* | |
| Bond lengths (Å) | 0.004 |
| Bond angles (°) | 0.605 |
| *Validation* | |
| Molprobity score | 1.84 |
| Clashscore | 7.68 |
| Poor rotamers (%) | 0.00 |
| Cβ outliers (%) | 0.00 |
| Cis-proline (%) | 7.50 |
| Cis-general (%) | 0.00 |
| Twisted proline (%) | 0.00 |
| Twisted general (%) | 0.00 |
| *Ramachandran plot* | |
| Favored (%) | 93.67 |
| Allowed (%) | 6.33 |
| Outliers (%) | 0.00 |

#### Table S2: Expression levels of mutant RABV-G constructs

Table reports expression level of each tested RABV-G construct. Median fluorescence intensity after staining with labelled RVC20 was measured, and is reported as a proportion of that observed with the appropriate wildtype (WT) comparator (i.e. untagged WT for untagged constructs, GFP-tagged WT for GFP-tagged constructs). Table reports median and range of four technical replicates across two experiments (a transfection with each of two independent DNA preparations on each of two days).

| Construct details | | | | | Expression (proportion of WT) | | |
|---|---|---|---|---|---|---|---|
| **Plasmid reference number** | **Residue number** | **Changed from** | **Changed to** | **Untagged or GFP fusion?** | **Median** | **Upper limit of range** | **Lower limit of range** |
| ADP502 | 20 | H | A | Untagged | 0.98 | 1.27 | 0.84 |
| ADP514 | 20 | H | L | GFP | 0.56 | 0.76 | 0.33 |
| ADP503 | 21 | H | A | GFP | 1.24 | 1.69 | 0.81 |
| ADP515 | 21 | H | L | GFP | 0.20 | 0.23 | 0.18 |
| ADP504 | 86 | H | A | Untagged | 0.08 | 0.15 | 0.06 |
| ADP516 | 86 | H | L | GFP | 0.17 | 0.34 | 0.13 |
| ADP505 | 113 | H | A | GFP | 0.84 | 1.53 | 0.20 |
| ADP517 | 113 | H | L | GFP | 0.72 | 2.24 | 0.35 |
| ADP506 | 150 | H | A | Untagged | 0.88 | 0.99 | 0.70 |
| ADP518 | 150 | H | L | GFP | 0.70 | 0.81 | 0.44 |
| ADP507 | 173 | H | A | GFP | 0.05 | 0.06 | 0.04 |
| ADP519 | 173 | H | L | Untagged | 0.02 | 0.02 | 0.02 |
| ADP508 | 261 | H | A | Untagged | 0.91 | 1.13 | 0.83 |
| ADP596 | 261 | H | L | Untagged | 1.01 | 1.07 | 0.75 |
| ADP520 | 261 | H | L | GFP | 0.80 | 1.04 | 0.72 |
| ADP600 | 266 | D | P | Untagged | 1.23 | 1.42 | 1.03 |
| ADP599 | 268 | I | P | Untagged | 0.87 | 1.09 | 0.74 |
| ADP484 | 269 | E | P | GFP | 0.04 | 0.05 | 0.03 |
| ADP593 | 270 | H | P | Untagged | 1.04 | 1.40 | 0.80 |
| ADP485 | 270 | H | P | GFP | 1.38 | 2.45 | 1.07 |
| ADP594 | 271 | L | P | Untagged | 1.10 | 2.02 | 0.90 |
| ADP486 | 271 | L | P | GFP | 1.05 | 1.69 | 0.82 |
| ADP595 | 272 | V | P | Untagged | 1.12 | 1.29 | 0.82 |
| ADP487 | 272 | V | P | GFP | 1.16 | 1.50 | 0.84 |
| ADP488 | 273 | V | P | GFP | 1.78 | 2.10 | 1.12 |
| ADP509 | 303 | H | A | GFP | 0.59 | 0.70 | 0.46 |
| ADP521 | 303 | H | L | Untagged | 0.93 | 1.22 | 0.79 |
| ADP510 | 328 | H | A | Untagged | 0.10 | 0.13 | 0.05 |
| ADP614 | 384 | H | K | Untagged | 0.91 | 1.22 | 0.62 |
| ADP601 | 384 | H | P | Untagged | 1.08 | 1.27 | 0.81 |
| ADP511 | 397 | H | A | Untagged | 0.09 | 0.17 | 0.06 |
| ADP523 | 397 | H | L | GFP | 0.05 | 0.06 | 0.04 |
| ADP512 | 419 | H | A | Untagged | 0.02 | 0.02 | 0.02 |
| ADP524 | 419 | H | L | Untagged | 1.03 | 1.31 | 0.67 |
| ADP513 | 424 | H | A | GFP | 0.44 | 0.90 | 0.14 |
| ADP525 | 424 | H | L | Untagged | 1.13 | 1.28 | 0.24 |
| ADP079 | WT | | | Untagged | 1.00 | 1.12 | 0.88 |
| ADP427 | WT | | | GFP | 1.00 | 1.17 | 0.83 |

#### References cited in Supplementary Data legends


Adams, P.D., Grosse-Kunstleve, R.W., Hung, L.W., Ioerger, T.R., McCoy, A.J., Moriarty, N.W., Read, R.J., Sacchettini, J.C., Sauter, N.K., and Terwilliger, T.C. (2002). PHENIX: building new software for automated crystallographic structure determination. Acta Crystallogr D Biol Crystallogr *58*, 1948-1954.

Corpet, F. (1988). Multiple sequence alignment with hierarchical clustering. Nucleic Acids Res *16*, 10881-10890.

Emsley, P., and Cowtan, K. (2004). Coot: model-building tools for molecular graphics. Acta Crystallogr D Biol Crystallogr *60*, 2126-2132.

Gouet, P., Robert, X., and Courcelle, E. (2003). ESPript/ENDscript: Extracting and rendering sequence and 3D information from atomic structures of proteins. Nucleic Acids Res *31*, 3320-3323.

Hellert, J., Buchrieser, J., Larrous, F., Minola, A., de Melo, G.D., Soriaga, L., England, P., Haouz, A., Telenti, A., Schwartz, O., *et al.* (2020). Structure of the prefusion-locking broadly neutralizing antibody RVC20 bound to the rabies virus glycoprotein. Nat Commun *11*, 596.

Kim, E.J., Jacobs, M.W., Ito-Cole, T., and Callaway, E.M. (2016). Improved Monosynaptic Neural Circuit Tracing Using Engineered Rabies Virus Glycoproteins. Cell Rep.

Krissinel, E., and Henrick, K. (2007). Inference of macromolecular assemblies from crystalline state. J Mol Biol *372*, 774-797.

Kuzmina, N.A., Kuzmin, I.V., Ellison, J.A., and Rupprecht, C.E. (2013). Conservation of Binding Epitopes for Monoclonal Antibodies on the Rabies Virus Glycoprotein. Journal of Antivirals & Antiretrovirals *5*, 37-43.

Laskowski, R.A., and Swindells, M.B. (2011). LigPlot+: multiple ligand-protein interaction diagrams for drug discovery. J Chem Inf Model *51*, 2778-2786.

Punjani, A., Rubinstein, J.L., Fleet, D.J., and Brubaker, M.A. (2017). cryoSPARC: algorithms for rapid unsupervised cryo-EM structure determination. Nature methods *14*, 290-296.

Roche, S., Rey, F.A., Gaudin, Y., and Bressanelli, S. (2007). Structure of the prefusion form of the vesicular stomatitis virus glycoprotein G. Science *315*, 843-848.

Yang, F., Lin, S., Ye, F., Yang, J., Qi, J., Chen, Z., Lin, X., Wang, J., Yue, D., Cheng, Y., *et al.* (2020). Structural Analysis of Rabies Virus Glycoprotein Reveals pH-Dependent Conformational Changes and Interactions with a Neutralizing Antibody. Cell host & microbe *27*, 441-453 e447.
