## Supplementary material for "Structure of trimeric pre-fusion rabies virus glycoprotein in complex with two protective antibodies": Materials and Methods

### Production of antibodies and Fabs

1112-1 IgG was produced from a hybridoma, which was a kind gift from Prof. Hindegund Ertl (Wistar Institute, USA), cultured using a CELLine CL 1000 Bioreactor (Cole-Parmer) and purified on a 1 mL Protein G column (ThermoFisher) as per the manufacturer's instructions. To prepare 1112-1 Fab for cryo-EM experiment, IgG was digested using a Mouse IgG1 Fab and F(ab')2 Preparation kit (ThermoFisher).

1112-1 heavy and light chain coding sequences were obtained by RNA sequencing (Absolute Antibody) and cloned into pVIP-ENTR, a mammalian expression vector modified from the pENTR4 vector (ThermoFisher) by incorporation of a synthetic 'CASI' promoter (a kind gift of Martino Bardelli) (Balazs et al., 2013). Briefly, coding sequences were synthesized (ThermoFisher) with appropriate adaptors for cloning using an In-Fusion Snap Assembly Kit (Takara Bioscience) and, for the heavy chain, C-terminal sequence encoding a TwinStrep tag (Schmidt et al., 2013). Synthetic genes were inserted into a linearized pVIP-ENTR vector using an In-Fusion Snap Assembly Kit (Takara Biosciences).

Sequences of the variable regions of RVC20, RVC58 and 17C7 have been disclosed (Corti, 2016; Hellert et al., 2020; Thomas et al., 2010). To produce the antibodies in Fab format, synthetic clonal genes carrying variable regions of the antibodies were synthesized by Twist Biosciences. Suitable adaptors were added to the genes to enable cloning into pOPINVH and pOPINVL vectors (Addgene #26041 and 26040) (Nettleship et al., 2008) using the In-Fusion Snap Assembly Kit. Heavy chain coding sequences included TwinStrep (TS) (RVC20, 17C7) or $His_6$ (RVC58) purification tags. pOPINVH was linearized with AfeI and XbaI restriction enzymes for RVC20-TS and 17C7-TS heavy chain cloning, or KpnI and SfoI for RVC58-His heavy chain cloning. pOPINVL was linearized with KpnI and SacI for cloning of all light chains.

To produce plasmids encoding RVC20 and 17C7 IgG heavy chains, VH regions for RVC20 and 17C7 were PCR-amplified from the above pOPINVH-based vectors. Forward and reverse PCR primers contained suitable InFusion adapters for further InFusion cloning of the PCR fragments in frame with a human IgG1 constant region coding sequence in the pVIP-ENTR backbone.

All recombinant Fabs and IgG were expressed by transient transfection of Expi293F cells (ThermoFisher), with a 1:1 heavy:light-chain-coding plasmid ratio, using the ExpiFectamine™ 293 Transfection Kit as per manufacturer's recommendations. Supernatants were harvested 96 hours after transfection and purified on either StrepTactin XT Superflow column (IBA Lifesciences) for RVC20-TS or 17C7-TS Fabs, Talon

Superflow column (Cytiva) for RVC58-His Fab, or Protein G column (ThermoFisher) for IgGs, as per manufacturer's recommendations.

### Production of RABV-G H270P

To produce untagged full-length RABV-G H270P-mutant protein, a stable cell line was generated essentially as previously described (Elegheert et al., 2018), with minor modifications.

The construct was based upon one previously described, composed of a codon optimized Pasteur strain ectodomain coding sequence chimerized with a SAD-B19 strain transmembrane and intracellular domain (Kim et al., 2016b), but encoding the desired H270P mutation. This was ligated into the pHR-CMV-TetO2-3C-Twin-Strep-IRES-Turquoise2 lentiviral shuttle plasmid (Addgene #113886), allowing production using a previously described method of a lentivirus encoding RABV-G H270P and mTurquoise fluorescent protein (the latter after an internal ribosome entry site), under the control of a tet-repressible promoter (Elegheert et al., 2018). HEK293-Trex cells (ThermoFisher) which had previously been adapted for growth in suspension in CD293 medium (ThermoFisher), were then transduced with this virus by spinoculation at 1300 x g for 2 h at 30 °C. Four days after transduction, RABV-G expression was partially induced using 0.01 µg/mL tetracycline and, 24 h later, the 5% of cells with the highest level of expression of mTurquoise were collected using an SH800 cell sorter (Sony). Sorted cells were expanded, initially in static culture in progressively larger vessels from a 24-well plate, in DMEM-F12 medium (ThermoFisher), supplemented with 10% fetal bovine serum (FBS), 1% penicillin/streptomycin and 1 µg/mL blasticidin (Melford Scientific). Cells were then returned to suspension culture in Erlenmeyer flasks, initially in CD293 medium (with 10% FBS, 4 mM GlutaMAX, 1% penicillin-streptomycin solution, and 1 µg/mL blasticidin), then, over two weeks, with progressive reduction of the amount of serum to 0.5% and replacement of CD293 medium with BalanCD293 medium (Fujifilm – Irvine Scientific).

For protein expression, cells were grown to $4x10^6$ cells/mL, diluted 50/50 with fresh BalanCD293 medium supplemented with 4 mM GlutaMAX, 1% Pen-Strep solution and 5% BalanCD feed (Fujifilm-Irvine Scientific), and induced with 10 µg/mL of tetracycline for 48 h. Cell pellets were collected, resuspended in storage buffer (50 mM HEPES, pH 7.5, 150 mM NaCl and 5% glycerol), and frozen at -80 °C.

To purify protein, thawed pellets were treated with extraction buffer (50 mM HEPES pH 7.5, 150 mM NaCl, 1% *n*-octyl-β-d-glucoside [Generon], cOmplete Protease Inhibitor cocktail [Roche]) supplemented with an excess of 17C7-TS Fab and Biolock solution (IBA Lifesciences) for 2 h at 4 °C, and the lysate was clarified by centrifugation at 4000 x g for 30 min. RABV-G – 17C7-TS complexes were purified from clarified lysate on a Strep-Tactin XT Superflow column (IBA Lifesciences) according to the manufacturer's recommendations, with wash and elution buffers supplemented with 1% *n*-octyl-β-d-glucoside. Fractions containing purified protein complex were mixed with 3-fold mass excess of A8-35 amphipol (Anatrace) and left at 4 °C for 1 h. Detergent was then removed using Pierce detergent removal spin columns (ThermoScientific), and the resulting sample was concentrated using an Amicon centrifugal concentrator (Millipore). An excess of 1112-1 Fab, and additional 17C7-TS Fab, were then added to saturate available binding sites, and the resulting RABV-G / 17C7-TS / 1112-1 complex was purified by size-exclusion chromatography on a Superose 6 10/300 Increase column (Cytiva) in buffer containing 50 mM HEPES pH 7.5 and 150 mM NaCl.

#### Cryo-EM grid preparation

Purified RABV-G / 17C7-TS / 1112-1 complexes were concentrated and mixed with *n*-octyl-β-d-glucoside to final concentrations of 1 mg/mL and 0.07%, respectively. 4 µL of the sample was pipetted on glow-discharged Quantifoil® holey carbon grids (1.2/1.3 µm copper 200 mesh) and was blotted for 3.5 s blot time at -15 N blotting pressure prior to flash freezing in liquid ethane using a Vitrobot Mark IV (FEI/ThermoFisher).

#### Cryo-EM data collection and processing and model building

Data acquisition was performed at the Electron Bio-Imaging Centre (eBIC) at Diamond Light Source (Harwell, UK). A 300 kV Titan Krios microscope (Thermo Fisher Scientific) equipped with a K3 direct electron detector (Gatan) and a 20 eV Gatan Imaging Filter (GIF) energy filter was used. Cryo-EM images were recorded automatically using SerialEM software in super resolution mode but saved with a 2x binning factor, at defocus values ranging from –0.8 to –2.6 µm at a calibrated magnification of 81,000x, corresponding to a pixel size of 1.06 Å/px. The total electron dose was 44.4 $e^{-}$/Å$^{2}$ over 45 frames (**Table S1**).

Recorded movies were motion corrected (global) and the contrast transfer function (CTF) parameters were determined using Gctf within cryoSPARC software (Punjani et al., 2017). Particles were automatically picked using circular blobs with a diameter of 80–150 Å. Picked particles were extracted from the micrographs with box sizes of 400 px and subsequently subjected to 2D classification. Per-particle motion correction (local) was performed on particles from selected 2D classes. *Ab-initio* reconstruction and 3D heterogeneous refinement were performed to generate 5 initial models and to classify the particles among them, respectively. Non-uniform refinement was performed on all 3D classes. Particles from the best model was used for another round of 3D heterogenous refinement (3 classes) to further classify for structural heterogeneity. The best models were selected for local (per-particle) CTF refinement to improve particle and map resolution and quality. C3 symmetry was applied for models selected for further reconstruction and refinement. Map resolution was estimated by gold-standard Fourier shell correlation (FSC) plot at the 0.143 threshold. Two maps at 2.8 Å and 3.2 Å were generated. The latter constitutes fuzzy densities around the membrane-proximal fusion domain and amphipol regions, therefore the 2.8 Å was selected for model building and refinement. The detailed data processing steps is summarized in a flowchart presented in **Figure S1**.

Model building for all protein chains was performed in COOT v.0.8.9.2 (Emsley and Cowtan, 2004) and refined by alternating cycles of real-space refinement in Phenix v.1.19.2 (Adams et al., 2002). Model geometry was validated using MolProbity v.4.5.1 (Chen et al., 2010). Geometry statistics, model B factors, and map vs model cross-correlation values are shown in **Table S1**. Map and model figures were generated using UCSF Chimera, ChimeraX (Pettersen et al., 2004; Pettersen et al., 2021), and PyMol. FSC curves and a local resolution histogram were plotted using data from cryoSPARC (Punjani et al., 2017), and correlation coefficient graphs were plotted using refinement data extracted from Phenix (Adams et al., 2002).

### Assessment of conformational stability of RABV-G by flow cytometry

In designing potentially stabilizing substitutions, we considered both the pre-fusion structure and the likely post-fusion structure, on the basis of low pH structures of RABV-G$^{ecto}$, VSV-G and Chandipura virus G (Baquero et al., 2015; Roche et al., 2006; Yang et al., 2020). Histidines have been suggested to act as pH sensors in lyssavirus glycoproteins, with protonation at low pH disrupting the pre-fusion structure. We therefore tested substitution of all histidines in the protein with alanine and leucine, with the following exceptions: for H270 and H384, substitution with proline was tested, as detailed below; for H328L, production of the expression construct failed; and H352/354 were not targeted as they do not mediate any notable interactions in the pre-fusion form and lie in regions near the protein surface not expected to be subject to rearrangement during conformational transition. We also introduced potentially helix-breaking proline substitutions at residues 266-272 and 384, which may rearrange from the L4 / L5 linkers into helices in the post-fusion protein. All evaluated substitutions are shown in Supplementary Table 2.

The same RABV-G coding sequence as was used for structural studies, but with desired point mutations (and, unless otherwise specified, encoding the wildtype H270), obtained as synthetic DNA (Twist Bioscience). In some cases, constructs were synthesized in frame with sequence encoding a C-terminal flexible linker, tobacco etch virus protease cleavage site, green fluorescent protein and a 6-His tag.

The complete encoded C-terminal tag sequence in these cases was:
GAGSAAGSGEFENLYFQGMVSKGEELFTGVVPILVELDGDVNGHKFSVSGEGEGDATYGKLTLKFICTTGKLPVPWPTLVTTLTYGVQCFSRYPDHMKQHDFFKSAMPEGYVQERTIFFKDDGNYKTRAEVKFEGDTLVNRIELKGIDFKEDGNILGHKLEYNYNSHNVYIMADKQKNGIKVNFKIRHNIEDGSVQLADHYQQNTPIGDGPVLLPDNHYLSTQSALSKDPNEKRDHMVLKEFVTAAGITLGMDELYKHHHHHH

The above constructs were digested with NotI and XbaI restriction enzymes, and ligated into a similarly digested pTT3 backbone (Durocher et al., 2002).

Expi293 cells were transfected with 1 µg/mL of plasmid encoding either full-length wild-type or mutant RABV G using ExpiFectamine™ reagent (ThermoScientific) as per the manufacturer's instructions. 300 µL of cell samples were harvested 48 h post-transfection, subjected to centrifugation at 300 x g for 5 min, and resuspended in phosphate buffered saline (PBS).

Binding of RVC20 IgG was used as an indicator of pre-fusion conformation (Supplementary Figure 11) (Hellert et al., 2020). 150 µL of the sample was washed in PBS (pH 7.3), whilst the other 150 µL was washed with an acidic buffer (50 mM Bis-Tris, 130 mM NaCl, pH 5.8). The samples were incubated in their respective buffer at room temperature for 30 mins. For RVC20, cell staining was performed with RVC20 IgG conjugated to Alexa Fluor 647 (Thermo Scientific), diluted in either PBS or acidic buffer to a concentration of 1 µg/mL, for 20 mins. For 1112-1 and 17C7, staining was performed similarly with IgG at 1 µg/mL, followed by secondary staining using Alexa Fluor 647-conjugated anti-mouse immunoglobulin secondary antibody (Thermo Scientific). Stained cells were subsequently washed and resuspended twice in their corresponding buffers before a final washing was repeated in PBS. Stained cells were sorted on a BD LSRFortessa™ Cell Analyzer (BD Biosciences). Using FlowJo v10 software (BD Biosciences) single cells were selected using forward scatter and side scatter-based gates, and histograms of fluorescence intensity for single cells were used to determine median fluorescence intensity (MFI) for each sample.

### Production of wild-type RABV-G for SPR

Surface plasmon resonance experiments used full length wildtype RABV-G with a C-terminal 'C-tag' (Glu-Pro-Glu-Ala). To make this, a construct was produced using a plasmid obtained from Addgene (#74288) encoding the Pasteur – SADB19 chimeric RABV-G described above (Kim et al., 2016a). The insert was amplified with primers F: TAGTAGGCGGCCGCCATGGTCCCACAGGCTCTCC and R: TAGTAGTCTAGATTTACGCTTCCGGTTCGAGCCGTGTCTCGCCCCC, followed by digestion with NotI and XbaI restriction enzymes, and ligation into a similarly digested pTT3 backbone (Durocher et al., 2002).

RABV-G C-tagged wildtype protein was transiently expressed in Expi293F cells (ThermoFisher) using the ExpiFectamine™ 293 Transfection Kit as per manufacturer's recommendations. 72 h after transfection, cell pellets were collected, resuspended in a storage buffer (50 mM HEPES, pH 7.5, 150 mM NaCl and 5% glycerol), and frozen at -80 °C.

To purify protein, pellets were treated with extraction buffer (50 mM HEPES pH 7.5, 150 mM NaCl, 1% n-octyl-β-d-glucoside, and Pierce™ Protease Inhibitor Mini Tablets, EDTA-free Thermo Scientific™ (ThermoFisher)) for 2 h at 4 °C, and the lysate was clarified by centrifugation at 4000 x g for 30 min. Following clarification, C-tagged RABV-G containing lysate was purified on a CaptureSelect™ C-tagXL affinity column (ThermoFisher) according to the manufacturer's recommendations with wash and elution buffers supplemented with 1% n-octyl-β-d-glucoside. After affinity purification, elution fractions were buffer-exchanged to the extraction buffer and once again subjected to C-tag-based affinity purification. Fractions containing purified RABV-G protein were then concentrated and subjected to size-exclusion chromatography on a Superose 6 10/300 Increase column (Cytiva) in a buffer containing 50 mM HEPES pH 7.5, 150 mM NaCl, and 1% n-octyl-β-d-glucoside.

### Surface plasmon resonance

Surface plasmon resonance (SPR) measurements were made using a Biacore S200 machine and software (Cytiva).

Neutral running buffer comprising 10 mM HEPES, 150 mM sodium chloride, 3 mM EDTA, and 0.05% polysorbate (HBS-EP+ ) was prepared and adjusted to pH 7.5. Acidic running buffer comprising 10 mM Bis-Tris, 150 mM sodium chloride, 3 mM EDTA, and 0.05% polysorbate (BBS-EP+ ) was prepared and adjusted to pH 5.6.

Single-cycle kinetics measurements of Fab – RABV-G interactions were performed using CM5 chips and amine coupling kit (both from Cytiva), at an analysis temperature of 4 °C. Attempted capture of RABV-G using a C-terminal affinity tag was unsuccessful. To provide a surface which could be regenerated with fresh RABV-G for each cycle, we therefore used a sandwich configuration, whereby C-tagged RABV-G was first captured by another (non-competing) anti-RABV-G IgG antibody to produce an active flow cell, prior to application of the Fab of interest. For measurement of RVC20 kinetics, capture was on 17C7 at both pH 7.5 and 5.6. For measurement of 1112-1 kinetics, capture was on RVC20 at pH 7.5, or on 17C7 at pH 5.6. For measurement of 17C7 kinetics, capture was on RVC20 at pH 7.5, or on 1112-1 at pH 5.6. Capture antibodies were immobilized at the maximum achievable densities (6000-12000 response unit, RU). An irrelevant IgG was immobilized at similar densities on a reference flow cell. At the start of each cycle, C-tagged wild-type RABV-G was applied to both active and reference cells, resulting in capture of approximately 100 RU on the active flow cells. Capture was presumed to be multi-valent and was stable, with dissociation of no more than 5.2% of bound RABV-G noted over 1650s (in control cycles in which buffer was used as analyte). For all three antibodies, recombinant Strep-tagged Fabs were used as analytes.

To collect an initial data set, at each of pH 5.6 and 7.5, one of the Fabs of interest was then injected at concentrations of 0.031, 0.125, 0.5, 2, 8, and 32 nM. No binding was observed for RVC20 at pH 5.6; in view of the presence of reduced binding for 1112-1 and 17C7 at pH 5.6, increased concentrations were subsequently used to improve data quality (1.15, 3.4, 10.3, 31, 93, 278 nM for 1112-1-TS Fab, and 0.27, 1.08, 4.34, 17.4, 69.5, 278 nM for 17C7-TS Fab). Each concentration of Fabs was injected for 240 s at a flow rate of 30 µL/min, followed by a final 1800 s dissociation phase. Data shown in Figure 4 is doubly-background-subtracted (i.e. after subtraction of responses on the irrelevant-IgG-coated reference flow cell and responses after injection of buffer). A 1:1 binding model was fitted using the Biacore S200 v2 Evaluation software. Extracted curves were plotted using Prism 9.0 (GraphPad software).

The ability of 17C7 and RVC58 to lock RABV-G in pre-fusion conformation was tested with similar equipment, materials and chip preparation, with the exceptions of use of an analysis temperature of 21 ˚C, use of a T200 instrument to produce the data shown in Figure 4 for 17C7, and preparation of an additional chip for which RVC58 was immobilized on the active flow cell. The injection series was as indicated on the Figure 4 legend. RVC20 Fab was injected at fixed 0.33 µM concentration.

### References cited in Methods section


Adams, P.D., Grosse-Kunstleve, R.W., Hung, L.W., Ioerger, T.R., McCoy, A.J., Moriarty, N.W., Read, R.J., Sacchettini, J.C., Sauter, N.K., and Terwilliger, T.C. (2002). PHENIX: building new software for automated crystallographic structure determination. Acta Crystallogr D Biol Crystallogr *58*, 1948-1954.

Balazs, A.B., Bloom, J.D., Hong, C.M., Rao, D.S., and Baltimore, D. (2013). Broad protection against influenza infection by vectored immunoprophylaxis in mice. Nature biotechnology *31*, 647-652.

Baquero, E., Albertini, A.A., Raux, H., Buonocore, L., Rose, J.K., Bressanelli, S., and Gaudin, Y. (2015). Structure of the low pH conformation of Chandipura virus G reveals important features in the evolution of the vesiculovirus glycoprotein. PLoS Pathog *11*, e1004756.

Chen, V.B., Arendall, W.B., 3rd, Headd, J.J., Keedy, D.A., Immormino, R.M., Kapral, G.J., Murray, L.W., Richardson, J.S., and Richardson, D.C. (2010). MolProbity: all-atom structure validation for macromolecular crystallography. Acta Crystallogr D Biol Crystallogr *66*, 12-21.

Corti, D. (2016). Antibodies that potently neutralize rabies virus and other lyssaviruses and uses thereof.

Durocher, Y., Perret, S., and Kamen, A. (2002). High-level and high-throughput recombinant protein production by transient transfection of suspension-growing human 293-EBNA1 cells. Nucleic Acids Res *30*, E9.

Elegheert, J., Behiels, E., Bishop, B., Scott, S., Woolley, R.E., Griffiths, S.C., Byrne, E.F.X., Chang, V.T., Stuart, D.I., Jones, E.Y., *et al.* (2018). Lentiviral transduction of mammalian cells for fast, scalable and high-level production of soluble and membrane proteins. Nat Protoc *13*, 2991-3017.

Emsley, P., and Cowtan, K. (2004). Coot: model-building tools for molecular graphics. Acta Crystallogr D Biol Crystallogr *60*, 2126-2132.

Hellert, J., Buchrieser, J., Larrous, F., Minola, A., de Melo, G.D., Soriaga, L., England, P., Haouz, A., Telenti, A., Schwartz, O., *et al.* (2020). Structure of the prefusion-locking broadly neutralizing antibody RVC20 bound to the rabies virus glycoprotein. Nat Commun *11*, 596.

Kim, E.J., Jacobs, M.W., Ito-Cole, T., and Callaway, E.M. (2016a). Improved Monosynaptic Neural Circuit Tracing Using Engineered Rabies Virus Glycoproteins. Cell Rep *15*, 692-699.

Kim, E.J., Jacobs, M.W., Ito-Cole, T., and Callaway, E.M. (2016b). Improved Monosynaptic Neural Circuit Tracing Using Engineered Rabies Virus Glycoproteins. Cell Rep.

Nettleship, J.E., Ren, J., Rahman, N., Berrow, N.S., Hatherley, D., Barclay, A.N., and Owens, R.J. (2008). A pipeline for the production of antibody fragments for structural studies using transient expression in HEK 293T cells. Protein Expr Purif *62*, 83-89.

Pettersen, E.F., Goddard, T.D., Huang, C.C., Couch, G.S., Greenblatt, D.M., Meng, E.C., and Ferrin, T.E. (2004). UCSF Chimera--a visualization system for exploratory research and analysis. J Comput Chem *25*, 1605-1612.

Pettersen, E.F., Goddard, T.D., Huang, C.C., Meng, E.C., Couch, G.S., Croll, T.I., Morris, J.H., and Ferrin, T.E. (2021). UCSF ChimeraX: Structure visualization for researchers, educators, and developers. Protein science : a publication of the Protein Society *30*, 70-82.

Punjani, A., Rubinstein, J.L., Fleet, D.J., and Brubaker, M.A. (2017). cryoSPARC: algorithms for rapid unsupervised cryo-EM structure determination. Nature methods *14*, 290-296.

Roche, S., Bressanelli, S., Rey, F.A., and Gaudin, Y. (2006). Crystal structure of the low-pH form of the vesicular stomatitis virus glycoprotein G. Science *313*, 187-191.

Schmidt, T.G.M., Batz, L., Bonet, L., Carl, U., Holzapfel, G., Kiem, K., Matulewicz, K., Niermeier, D., Schuchardt, I., and Stanar, K. (2013). Development of the Twin-Strep-tag® and its application for purification of recombinant proteins from cell culture supernatants. Protein Expression and Purification *92*, 54-61.

Thomas, W.D., Jr., Ambrosino, D.M., Mandell, R.B., Sloan, S.E., Babcock, G.J., and Rupprecht, C.E. (2010). Human Antibodies Against Rabies and Uses Thereof.

Yang, F., Lin, S., Ye, F., Yang, J., Qi, J., Chen, Z., Lin, X., Wang, J., Yue, D., Cheng, Y.*, et al.* (2020). Structural Analysis of Rabies Virus Glycoprotein Reveals pH-Dependent Conformational Changes and Interactions with a Neutralizing Antibody. Cell host & microbe *27*, 441-453 e447.
